# Structural origins of *Escherichia coli* RNA polymerase open promoter complex stability

**DOI:** 10.1101/2021.09.08.459427

**Authors:** Ruth M. Saecker, James Chen, Courtney E. Chiu, Brandon Malone, Johanna Sotiris, Mark Ebrahim, Laura Y. Yen, Edward T. Eng, Seth A. Darst

## Abstract

The first step of gene expression in all organisms requires opening the DNA duplex to expose one strand for templated RNA synthesis. In *Escherichia coli*, promoter DNA sequence fundamentally determines how fast the RNA polymerase (RNAP) forms “open” complexes (RPo), whether RPo persists for seconds or hours, and how quickly RNAP transitions from initiation to elongation. These rates control promoter strength *in vivo* but their structural origins remain largely unknown. Here we use cryo-electron microscopy to determine structures of RPo formed *de novo* at three promoters with widely differing lifetimes at 37°C: λP_R_ (t_1/2_ ∼ 10 hours), T7A1 (t_1/2_ ∼ 4 minutes), and a point mutant in λP_R_ (λP_R-5C_) (t_1/2_ ∼ 2 hours). Two distinct RPo conformers are populated at λP_R_, likely representing productive and unproductive forms of RPo observed in solution studies. We find that changes in the sequence and length of DNA in the transcription bubble just upstream of the start site (+1) globally alter the network of DNA-RNAP interactions, base stacking, and strand order in the single-stranded DNA of the transcription bubble; these differences propagate beyond the bubble to upstream and downstream DNA. After expanding the transcription bubble by one base (T7A1), the nontemplate-strand “scrunches” inside the active site cleft; the template-strand bulges outside the cleft at the upstream edge of the bubble. The structures illustrate how limited sequence changes trigger global alterations in the transcription bubble that modulate RPo lifetime and affect the subsequent steps of the transcription cycle.

## Introduction

Transcription by DNA-dependent RNA polymerases (RNAP) releases information stored in duplex DNA in the form of RNA transcripts. Appropriate responses to changing cellular conditions and growth rates require rapid and tight control of cellular RNA levels. In the model organism *Escherichia coli*, the rate of productive initiation events largely determines RNA transcript amount (1). RNA chain initiation frequencies vary over four orders of magnitude *in vivo*; during exponential growth 10^3^-10^4^ ribosomal transcripts are synthesized per generation whereas other transcripts may appear once or not at all (1-3). Intensive investigations of how *E. coli* achieves this extraordinary range are ongoing, recently revealing novel modes of transcription initiation (4) and termination (5, 6) despite decades of study.

In bacteria, a single “core” enzyme (E), comprising five subunits (α_2_ββ’ω), catalyzes all templated phosphodiester bond synthesis. Operon-specific output is orchestrated by the addition of a sixth dissociable subunit sigma (σ), forming the holoenzyme (Eσ) [cf. (3, 7)]. The vast majority of studies have focused on the “housekeeping” group I σ factors [σ^70^ in *Eco*; σ^A^ in other bacteria (8)]. Regions of high sequence conservation in the σ^70^-family (numbered sequentially 1.1-4.2) correspond to structural domains linked by flexible linkers (9, 10), each playing distinct roles in promoter DNA recognition and strand separation [cf. (3)].

Exposing and positioning the start site (+1) near the RNAP catalytic Mg^2+^ in the RNAP open promoter complex (RPo) requires unwinding of over a turn of the DNA helix (11). Because binding free energy drives these steps, promoter DNA sequence intrinsically determines how quickly RPo forms and how long it persists (1, 12-14). These kinetic differences critically underlie cell function. For example, differences in RPo lifetime allow the RNAP-binding factors dksA and ppGpp to “discriminate” between the operons they regulate and those they do not, and to have opposing effects at those they do (15, 16).

A striking discovery of ensemble and single molecule mechanistic investigations is that RPo is not a singular universal complex; multiple forms of RPo can exist at the same promoter (cf. (17-22)]. At the model phage promoter λP_R_, a series of distinct open complexes form after the rate-limiting step [I1 to I2; (21, 22)]:

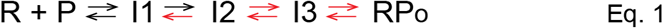

[Eq. 1. Minimal mechanism of RPo formation at λP_R_. Steps that contribute to the observed dissociation rate constant shown in red (11, 21)]. I2 is only transiently populated; conversions to I3 and RPo successively stabilize the strand-separated state. The structural and functional differences between these complexes are largely unknown.

While the conserved -35 and -10 elements and the length of the “spacer” between them affect RPo formation and thus its overall stability, both the sequence and length of the “discriminator” [Fig. 1*A*; (16)] has emerged as a primary determinant of RPo lifetime (23-25). Haugen et al. (23) demonstrated that the critical sequence runs from -6 to -4 (numbering with respect to the transcription start site at +1), with 5’-GGG-3’ [nontemplate strand (nt-strand) sequence] yielding the longest RPo half-lives (t_1/2_)(23). The cornerstone of “discrimination” was pinpointed to -5 on the nt-strand, where the presence of G [G_-5_(nt)] increases RPo t_1/2_ by 10 to 50-fold at the *rrnB* P1, λP_R_, λP_L_ and P*gal* promoters. Crosslinking, mutational and other biochemical approaches mapped these large effects on RPo stability to specific interactions with σ^70^ conserved region 1.2 (σ^70^_1.2_) (26, 27). Subsequent work found that the β “gate loop” (βGL) also plays a crucial role in nt-strand discriminator interactions (28). How do these interactions dictate RPo lifetimes that range from seconds to many hours?

**Fig. 1.**
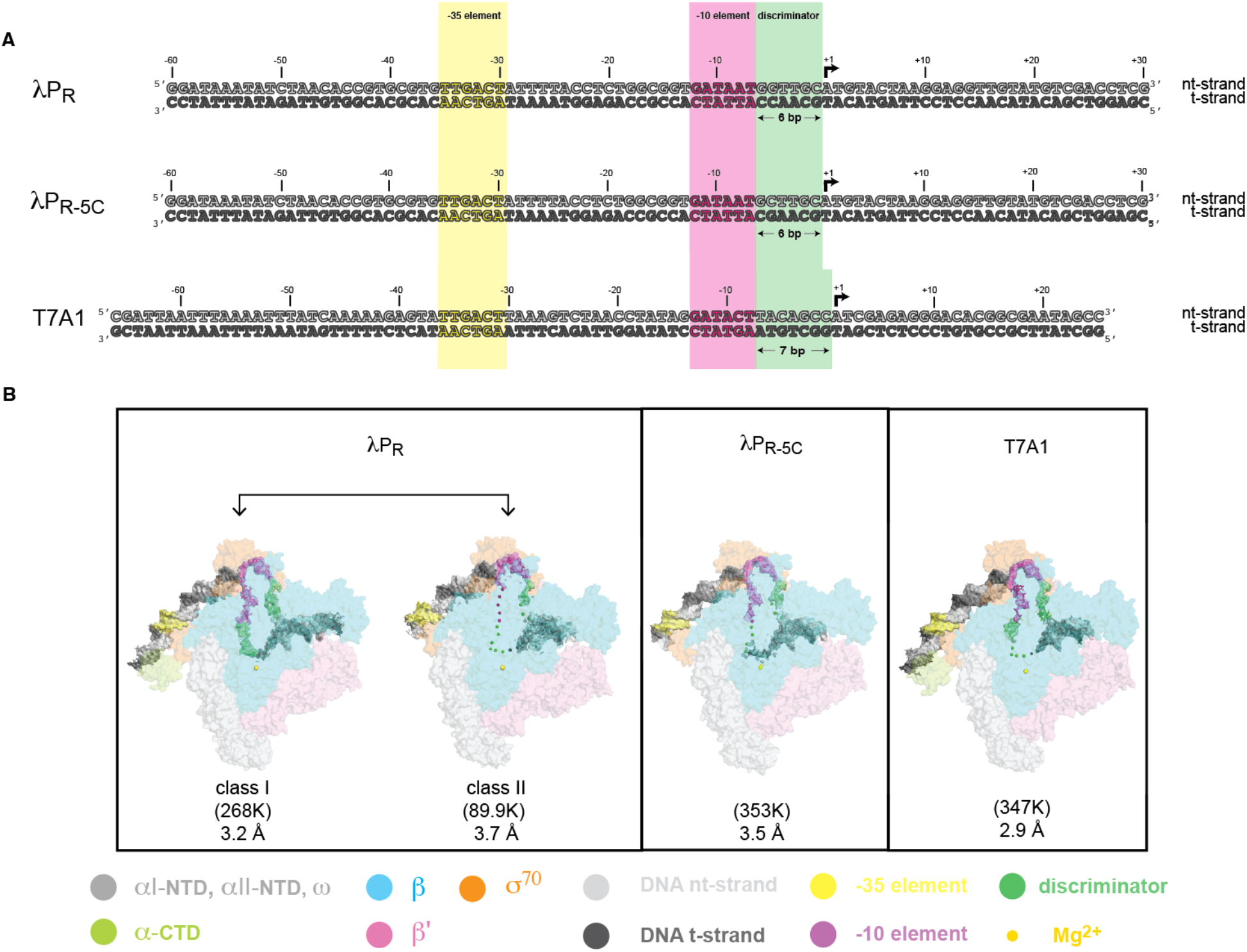
Promoter DNA constructs used for cryo-EM studies and overall cryo-EM structures of RPos. (*A*) Promoter sequences studied by cryo-EM [nt-strand DNA (top strand), light grey; t-strand (bottom strand), dark grey]. Numbers above the DNA sequences denote positions with respect to the transcription start site (+1, denoted by the black arrow). Shaded colors highlight key promoter regions: -35 element (yellow), -10 element (magenta), and the discriminator (pale green). (*top*) λP_R_ (−60 to +30). -60 to +20 are native λP_R_ sequences. Sequences downstream of +20 originate from the plasmid construct used in extensive kinetic and DNA footprinting studies of λP_R_ (11); (*middle*) λP_R-5C_ (−60 to +30). Single base pair inversion of λP_R_ G_-5_ to C. (*bottom*) T7A1 (−66 to +20). While T7A1 and λP_R_ share the same -35 element sequence, key differences exist in both the -10 element and the sequence and length (7 vs 6 nucleotides, respectively) of the discriminator. At the upstream end of the constructs used here, the T7A1 promoter has both proximal and distal UP elements (tight-binding αCTD binding sites), and λP_R_ (and λP_R-5C_) has a distal UP (3). (*B*) *Eco* RNAP subunits are shown as transparent surfaces (αI, αII, w, light gray; αCTD, pale green; β, pale cyan; β’, light pink; σ^70^, light orange) with the active site Mg^2+^ shown as a sphere (pale yellow). Promoter DNA is shown as cryo-EM difference density (−35 element, -10 element and discriminator colored as in *A*). Particle classification revealed two distinct RNAP-DNA complexes populated at λP_R_, and a single class at T7A1 and at λP_R-5C_. The number of particles in each class and nominal resolution are shown. In λP_R_ class I, good map density allowed all bases in the nt- and t-strands to be modeled. For comparison, disordered bases in the other RPo are shown as spheres positioned approximately at the corresponding position of the phosphate backbone in λP_R_ class I. DNA in RPo modeled from -45 to +23 in λP_R_ class I, -37 to +13 in λP_R_ class II, -37 to +15 in λP_R-5C_, and -47 to +15 in T7A1.

The understanding of how the transcription bubble is differentially stabilized is currently limited. The first atomic resolution structures of RPo determined by X-ray crystallography used promoters with consensus -35 and -10 promoter elements and pre-formed bubble templates with short regions of downstream duplex (29, 30) or short downstream “fork” constructs (31, 32). Resulting structures often revealed strand disorder; to improve resolution, NTPs or short RNA primers were added, forming transcription initiation complexes (RPinit). Until very recently (33-36), the structure of RPo was mostly inferred from these RPinit’s formed at nonnative sequences/structures.

Similarly, detailed biochemical studies of RPo formation exist for only a small handful of promoters. Mechanistic and biochemical investigations of DNA opening at two phage promoters (λP_R_ and T7A1) have largely defined the critical steps in forming the transcription bubble [cf. (11, 14, 37-39)]. However, no high resolution structural data exist for any complex formed at either promoter. Here we begin to address these gaps by using cryo-electron microscopy (cryo-EM) to visualize RPo formed *de novo* at λP_R_ and T7A1. We also studied a single point mutant in λP_R_ (λP_R-5C_) that significantly decreases RPo half-life. Below we present the structural differences between these complexes, providing insights into how changes in promoter sequence, even at a single base, give rise to orders of magnitude changes in RPo lifetimes.

## Results

### Cryo-EM structures of Eα^70^ RP_o_ at the λP_R_, λ_PR-5C_, and T7A1 promoters

To understand how promoter sequence dictates widely differing RPo lifetimes, we analyzed three *de novo* DNA-melted Eσ^70^-promoter complexes by single particle cryo-EM [T7A1, λP_R_, and a point mutant at -5 in λP_R_ (λP_R-5C_; Fig. 1*A*). To reduce particle orientation bias at the liquid-air interface, the cryo-EM buffer contained 8 mM CHAPSO (40). CHAPSO decreases RPo lifetime (2 to 3-fold) but does not change the relative stabilities of each RPo (SI Appendix, Fig. S1 and Table S1). Steps of maximum likelihood classification (41) revealed two distinct conformational classes populated at λP_R_ (75% of the particles fall into class I, 25% in class II) whereas only a single class was found for T7A1 and λP_R-5C_ (Fig. 1*B*; SI Appendix, Figs. S2-S8 and Table S2).

In all complexes: i. the transcription bubble is fully open; and ii. interactions with the -35 element, the spacer region and the -10 element bases on the nt-strand are largely the same. Because these interfaces do not significantly differ from each other or from previously reported structures [cf. (10, 29-31, 34, 42)] they will not be discussed further.

Relative to λP_R_ class I, the DNA strands in transcription bubbles of the other RPo are more dynamic. Every single base in the nt- and t-strands within the transcription bubble is well-resolved in λP_R_ class I (Figs. 1*B* and 2*A*) whereas the entire t-strand (−11 to +2) and -4 to +2 on the nt-strand have little to no map density in class II (Figs. 1*B* and 2*B*). Similarly, the mid-regions of the λP_R-5C_ bubble (Figs. 1*B* and 2*C*) and the T7A1 t-strand from -4 to +2 (Figs. 1*B* and 2*D*) could not be modeled.

Duplex DNA regions distant from the transcription bubble also exhibit distinct differences. Unlike the relatively unstable RPos (SI Appendix, Fig. S1 and Table S1), an additional helical turn could be modeled downstream of +13 (to +23) in λP_R_ class I. Upstream of the -35 element, map density for the two flexibly-tethered αCTDs bound to DNA (−38 to -55) exists for all RPo but only the “proximal” αCTD in λP_R_ class I and T7A1 RPo was well-resolved [Fig. 2 and SI Appendix, Figs. S3 and S8, respectively; see also (43, 44)]. “Distal” αCTD (UP) sequences exist at both promoters (3), prompting a focused classification of this region. This approach extracted classes with different DNA trajectories but not distinct bound states of the second αCTD (cf. (35)). Thus despite the presence of UP elements in these promoters and in the ribosomal promoters *rpsT* P2 and *rrnB* P1, by the time RPo has formed, the DNA upstream of -45 is dynamic and the DNA binding mode of the second αCTD is heterogeneous (34-36, 45).

**Fig. 2.**
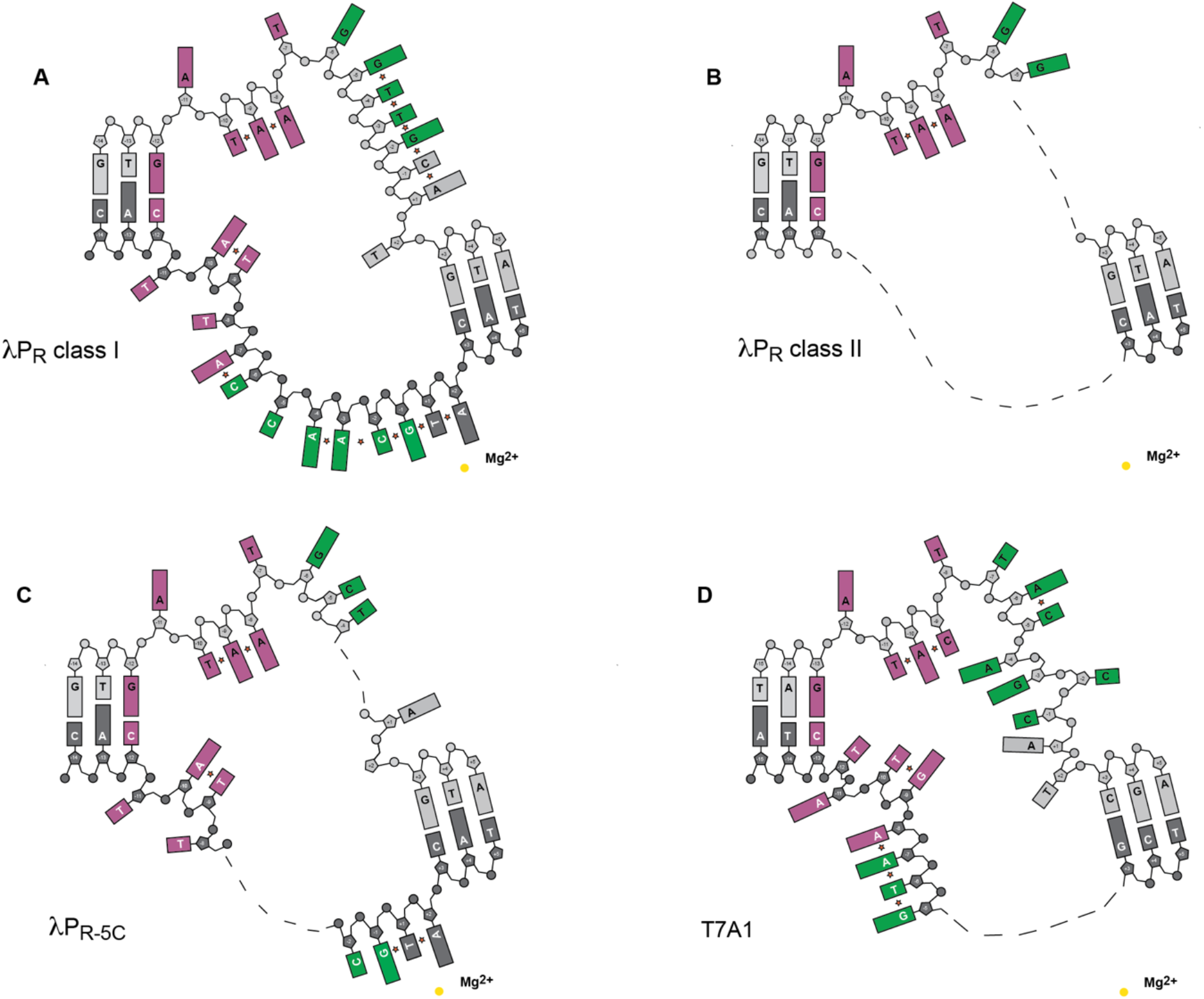
Comparison of differences in base stacking in the transcription bubble. The schematic illustrates: i. position of bases in each open complex, missing bases shown as dashes; and ii. base stacking pairs indicated by a star symbol. DNA backbone and bases colored as in Fig. 1. (*A*) λP_R_ class I. (*B*) λP_R_ class II. (*C*) λP_R-5C_. (*D*) T7A1.

Superposition of the cryo-EM RPo structures here and published high resolution cryo-EM RPo complexes [*rpsT* P2 (34) and *rrnB* P1 (36)] revealed small to moderate differences in the conformation of the clamp and the βlobe (SI Appendix, Table S3). With respect to λP_R_ class I, the clamp and the βlobe are more open in all other RPo. Additional narrowing of the cleft in λP_R_ class I appears to be stabilized by partial ordering of five σ^70^ residues (σ^70^S85 to S89; SI Appendix, Fig. S9) on the βlobe. Extension across the βlobe creates a “clasp” between β’ and β similar to that observed in mycobacterial Eσ^A^ (33); the clasp must be undone for DNA to leave the cleft.

σ^70^S85 to S89 lie at the C-terminus of the 37-residue linker connecting σ^70^_1.2_ to σ^70^_1.1_ [σ^70^_1.1_ -linker; (46)]. During RPo formation, DNA displaces σ^70^_1.1_ bound in the cleft [cf. (35, 47)]. No density exists for σ^70^_1.1_ in any RPo structure to date, indicating it does not rebind elsewhere. Instead, ejection creates a high local concentration of the flexibly-tethered σ^70^_1.1_ above the cleft. Interactions between the linker and the βlobe may increase stability in λP_R_ class I by directing σ^70^_1.1_ away from the channel, effectively reducing its concentration near the downstream DNA and disfavoring its re-entry relative to other RPo (see SI Appendix for a discussion of relationship of class I and class II to the intermediates in Eq. 1).

### Differences in base stacking in the transcription bubble

To illustrate the differences in strand/base resolution and in the extent of base stacking, each transcription bubble is presented schematically in Fig. 2. At the upstream end, the nt-strand single-stranded -10 element hexamer exhibits the same conserved interactions seen in all RPo structures to date: A_-11_(nt) and T_-7_(nt) are flipped out into protein pockets on σ, the intervening bases from -10 to -8 stack with each other, facing into the channel with only the backbone atoms making interactions with σ. However, downstream of the -10 element, differences in the discriminator length and sequence impact both single-stranded base stacking and strand order. For λP_R_ class I, almost every base in the bubble (nt- and t-strands) has a stacking partner (Fig. 2*A*). The single base change from G to C (nt-strand) at -5 of λP_R-5C_ affects the entire bubble, disordering regions of both strands (Fig. 2*C*). Addition of an additional nucleotide to the discriminator (T7A1) and placing an A at the same position as G_-5_(nt) in λP_R_ [A_-6_(nt) T7A1 numbering] disrupts all base stacking interactions in the nt-strand downstream of -5 (Fig. 2*D*).

### Interactions with the nt-strand: structural consequences of base identity at -5 and of a 7 base vs 6 base discriminator

How do three nt-strand bases out of all the bases that define a promoter profoundly effect RPo lifetime (23)? DNA opening positions bases from -6 to -4 near highly conserved residues in σ^70^_1.2_, σ^70^_2_ (9, 10) and the opposing βGL [cf. (31, 48, 49)]. These interactions close off the top of the active site channel, effectively trapping the strand-separated DNA inside (see SI Appendix, Figs. S3*C*, S4*C*, S6*C* and S8*C*). As detailed below, changes away from guanine (G) and increasing discriminator length significantly reduce the extent of these contacts, resulting in global changes in RPo structure.

At λP_R_, base stacking and pairing interactions from -6 to -4 in the duplex DNA are replaced in RPo by an extensive network of polar, π, and van der Waals contacts with highly conserved Eσ^70^ residues (Fig. 3*A*). Notably, σ^70^M102, σ^70^R99, and βR371 interact with multiple bases in λP_R_ class I RPo but not in T7A1 (Fig. 3*B*) or λP_R-5C_ (SI Appendix, Fig. S11). The “keystone” G_-5_(nt) (23) is tightly held by distinct chemical interactions that include two base-specific hydrogen bonds [σ^70^R99(NE)-O6 and σ^70^D96(OD2)-N1; Fig. 3*A*]. Replacing λP_R_ G_-5_(nt) with C not only eliminates these local contacts but abolishes the complex set of interactions that constrain the bases and sugar phosphate backbone from -6 to -4 in λP_R_ (SI Appendix, Fig. S11). The quality and quantity of interactions in the T7A1 nt-strand interface are diminished as well (Fig. 3*B*). Because Eσ^70^ intimately reads out the bases at -6 to -4 (Fig. 3*A*), deviations from the optimal sequence (G) as well as length alter the interface cooperatively. As a result, the driving force for the isomerizations that stabilize RPo changes nonadditively with sequence, yielding widespread differences in RPo structure (the extent of strand and downstream DNA order, clamp/βlobe/σ^70^_1.1_ -linker positions; Fig. 3, SI Appendix, Figs. S10 and S11, Table S3) and lifetime.

**Fig. 3.**
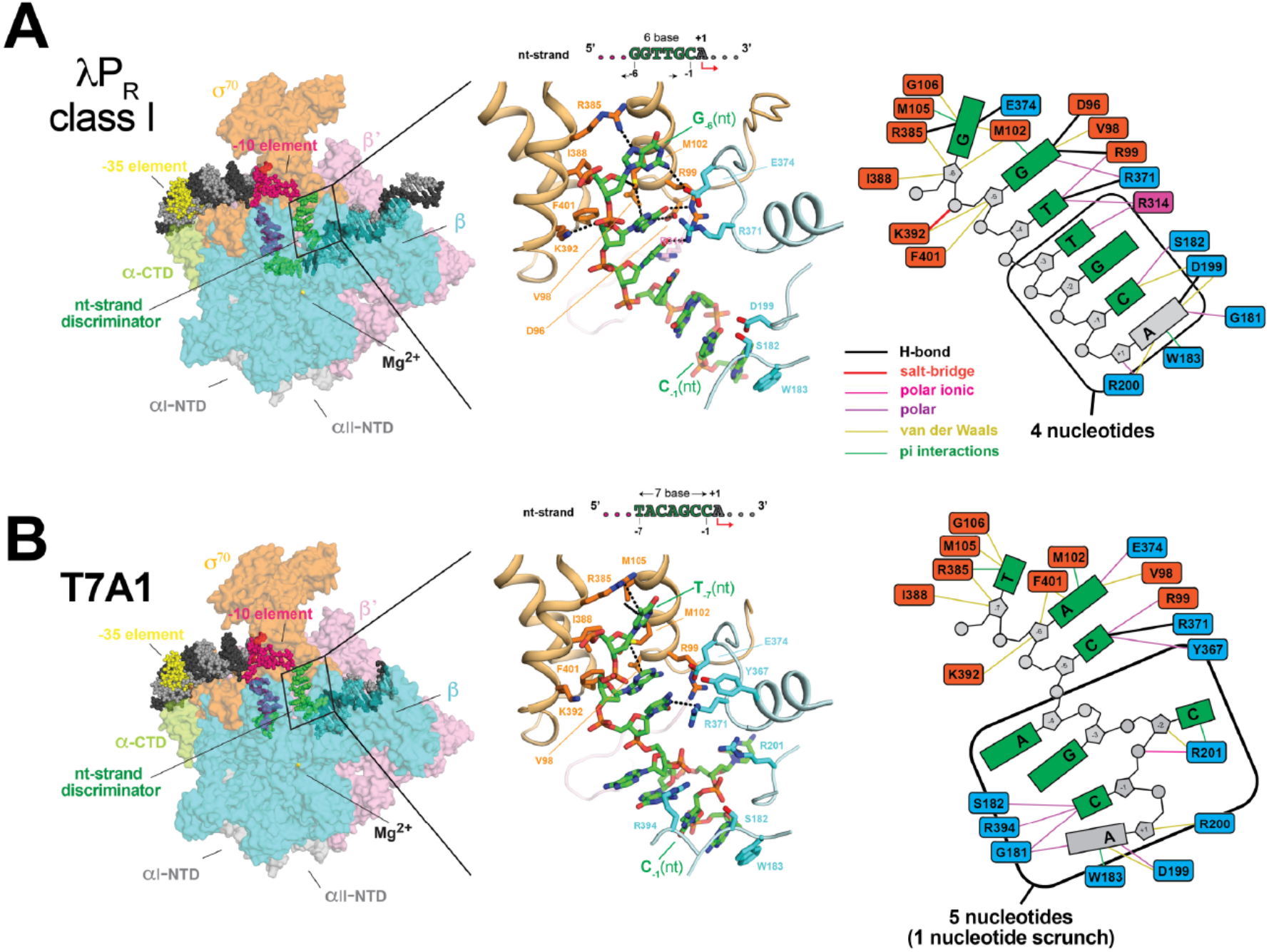
Differences in Eσ^70^ interactions with the nt-strand discriminator region between *(A)* λP_R_ class I (6 base discriminator) and (*B*) T7A1 (7 base discriminator). (*Left*) Overall view of RPo (similar to Fig. 1*B* with same color scheme) with RNAP subunits shown in surface representation and DNA as atomic spheres. The boxed area is magnified in the middle. (*Middle*) Magnified view showing interactions with the nt-strand from the discriminator to +1. RNAP subunits shown in backbone worm (β, pale cyan; β’, light pink; σ^70^, light orange)); sidechains of atoms within 4.5 Å of nucleic acid atoms [λP_R_(−6 to +1) or T7A1(−7 to +1)] are shown as sticks. Atomic distances within 3.5 Å and interactions with the π electrons of the DNA bases (sulfur-π, and cation-π) are shown by black dashed lines. The DNA carbon atoms are colored green. (*Right*) Schematic comparing the network of interactions between the nt-discriminator (and +1) and Eσ^70^. Favorable interactions within 3.5 Å (hydrogen bonds, salt bridges) are shown by heavier lines (see color key) than those within 4.5 Å [polar, ionic, van der Waals, π (sulfur, cation, π)]. Corresponding cryo-EM density is shown in SI Appendix, Fig. S10. Although the nature and extent of interactions with the nt-strand discriminator differ between λP_R_ and T7A1, the DNA backbone is largely solvent-exposed. Few favorable interactions exist with the sugar atoms (at upstream and downstream ends of the discriminator region) and only one positively charged amino acid falls within 4.5 Å of the DNA phosphate backbone of either promoter.

One key similarity unites all RPo characterized to date: the majority of the nt-strand discriminator contacts are with the upstream and downstream ends, leaving the mid-region fairly unconstrained and indeed observed to be dynamic in λP_R-5C_ and in λP_R_ class II. As seen in T7A1, adding another base to the canonical six base discriminator is accommodated by unstacking and flipping bases out of the backbone relative to one another in the “middle” of the discriminator (Fig. 3*B*). In this “scrunch”, A_-4_(nt) and G_-3_(nt) face into the RNAP channel where they are solvent-exposed, and C_-2_(nt) occupies the downstream end of the channel where it interacts with βR201.

### DNA scrunches in the t-strand at the upstream end of the DNA channel in RPo

Entry of the t-strand into the active site channel at T_-11_(t) [T_-12_(t) in T7A1] distorts the DNA backbone, unstacking and flipping T_-11_(t) out at λP_R_ and both T_-12_(t) and A_-11_(t) at T7A1, placing the t-strand “scrunch” at the upstream end of the channel (Fig. 4). While map density exists for these bases in locally filtered maps, it is weaker than that for the corresponding base partners on the nt-stand, consistent with the accessibility of these conserved thymines to permanganate ions (20, 50). As the strand descends further into the cleft, both λP_R_ and T7A1 exhibit an intriguing interaction with the single-stranded -10 element t-strand at A_-10_(t)/T_-9_(t) or T_-10_(t)/G_-9_(t), respectively. In both RPo, these bases are stacked and captured by the loop between the α-helices formed by σ^70^_2.1_ and σ^70^_2.2_, and the opposing residues on the βprotrusion (Fig. 4, SI Appendix, Fig. S12). At the upstream end of this “sandwich”, σ^70^R397 and σ^70^N396 interact with the -10 base, the -10 phosphate oxygen makes polar ionic contacts with σ^70^R468 (σ^70^_3_ helix). At the downstream end, the -9 base and βR470 form a cation–π stacking interaction and βK496 makes both ionic and nonpolar interactions with the DNA backbone (Fig. 4). Whether the “sandwich” causes the scrunch or whether it helps stabilize the scrunch is unknown. However, placing the additional base (relative to 6 base discriminator) at the upstream end of the channel equalizes the length of the t-strand from -8 to the active site (see Discussion).

**Fig. 4.**
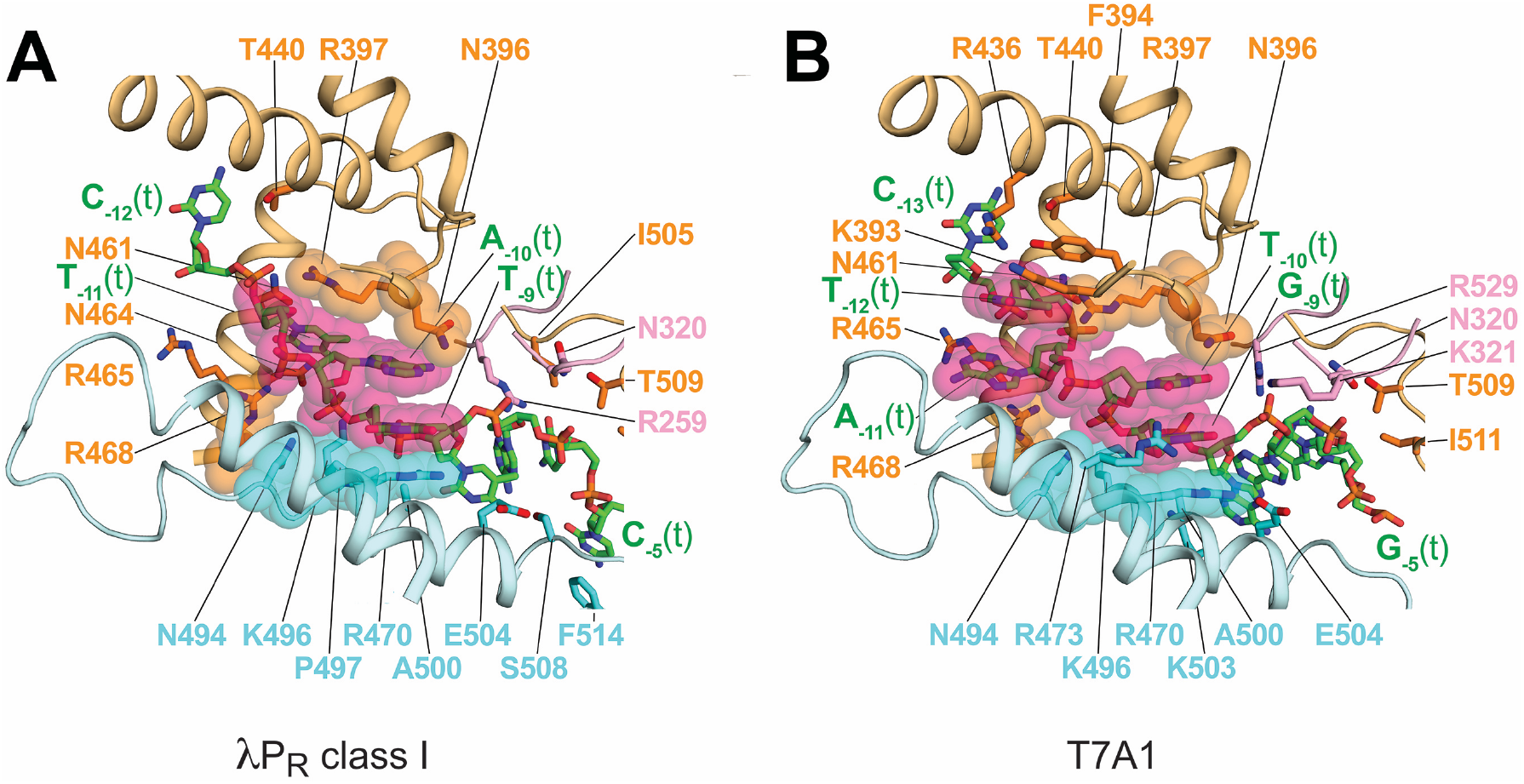
Scrunching in the t-strand. (*A*) λP_R_ class I RPo flips out T_-11_(t) at the ss/ds junction. (*B*) T7A1 flips out two bases, T_-12_(t) and A_-11_(t), scrunching the t-strand at the upstream entrance to the active site channel. Residues within 4.5 Å of nucleic acid atoms of the t-strand are shown as sticks. *Eco* Eσ^70^ subunits shown in backbone worm (β, pale cyan; β’, light pink; σ^70^, light orange). Single-stranded bases from -11 to -9 (*A*; λP_R_) or -12 to -9 (*B*; T7A1) are shown as sticks (same color coding as Fig. 3) and also transparent atomic spheres (hot pink). The same σ and β residues stabilize the A_-10_(t)/T_-9_(t) or T_-10_(t)/G_-9_(t) stacking pairs in λP_R_ and T7A1, respectively, noted and shown as transparent CPK atoms (σ, orange; β, cyan). Corresponding cryo-EM density is shown in SI Appendix, Fig. S12.

### λP_R_ t-strand is largely stacked and tightly held in the active site channel

The relatively high resolution of all the transcription bubble bases of the λP_R_ class I RPo and of the residues in the active site channel allows a molecular visualization of the interactions that direct the t-strand from the ds/ss upstream fork to the Mg^2+^ active site on the “floor” of the cleft some 65 Å away. As illustrated in the stick/cartoon representation (Fig. 5*A*) and schematic (Fig. 5*B*), every base in the upstream half of the bubble makes multiple favorable interactions with at least one residue of σ^70^, β and/or β’. While flipped out of the helix at the ds/ss junction, T_-11_(t) is not captured in a protein pocket like its base pairing partner on the nt-strand. Nonetheless, σ^70^ appears to constrain its position via numerous interactions with the base and backbone atoms: i. polar contacts between thymine and a trio of residues in the σ^70^_3_ helix, including a H-bond between σ^70^N461 and N3; ii. salt-bridges between the phosphate oxygens of C_-12_(t) and T_-11_(t) and σ^70^R397 and σ^70^R468, respectively; and iii. multiple polar and nonpolar contacts with sugar atoms. With the exception of σ^70^N461, these residues are invariant or nearly invariant in the housekeeping σ factors. Multiple interactions form between Eσ^70^ and the bases and the sugar phosphate backbone from -11 to -6, including H-bonds with T_-11_(t), A_-7_(t), and C_-6_(t). As seen for the nt-strand, π interactions form with unstacked bases at T_-9_(t) (βR470), C_-5_(t) (σ^70^F514), and A_-4_(t) (σ^70^F522).

**Fig. 5.**
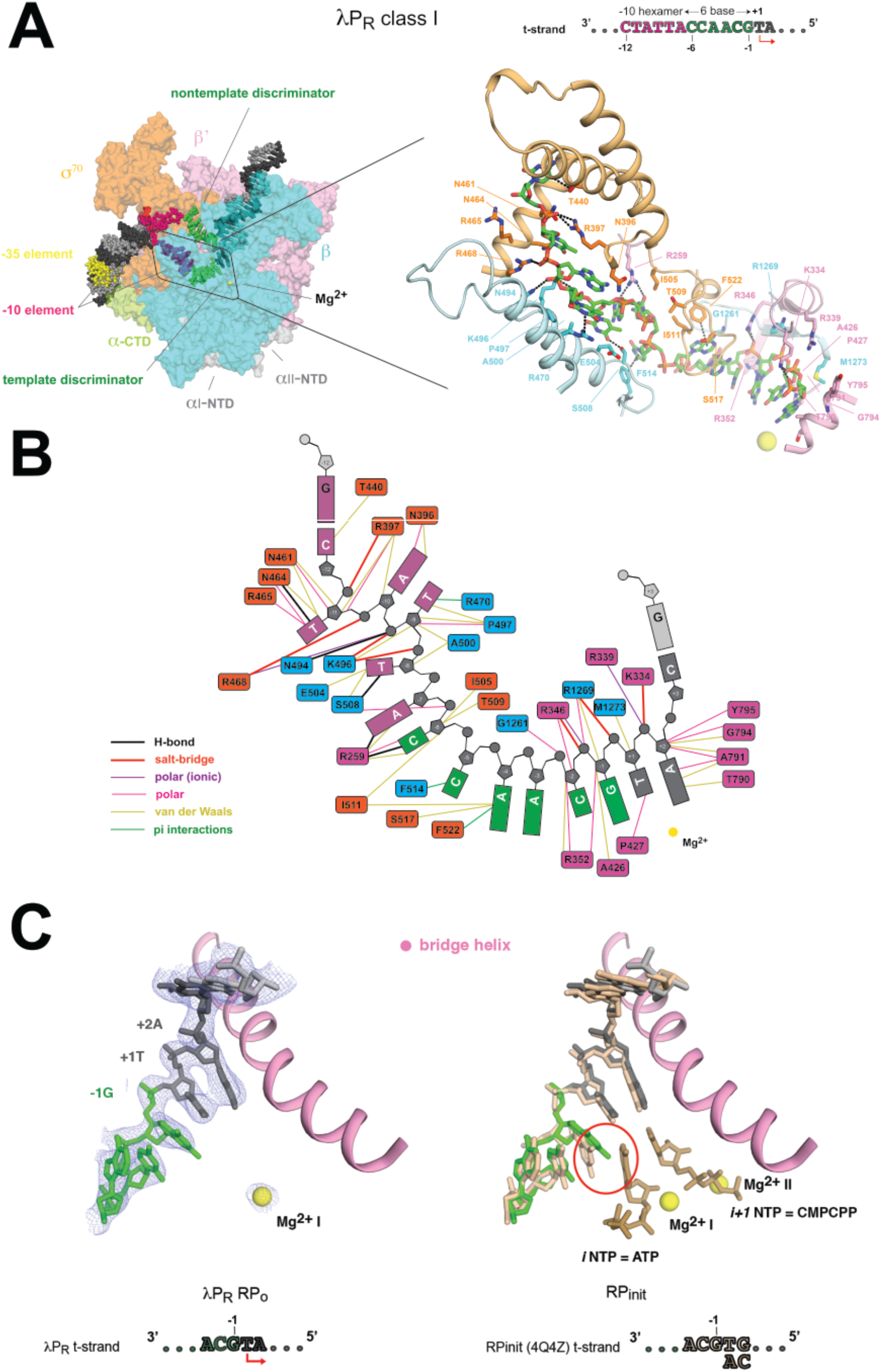
λP_R_ class I t-strand interactions and positioning in the active site. (*A*) (*Left*) Overall representation of RPo as in Fig. 3. The boxed area is magnified to the *right*. (*B*) Schematic detailing the interactions between the t-strand and residues within 4.5 Å. (*C*) Comparison of the t-strand in λP_R_ class I RPo and in RPinit. (*Left*) Cryo-EM density (blue mesh) defining the modeled position of single-strand bases on the t-strand (−3 to +2), the downstream ss/ds junction at +3 (shown as sticks) and the Mg^2+^ bound in the RPo active site (yellow sphere). The conserved RNAP bridge helix (β’; pink) is shown as a point of reference. Colors as in Fig. 1. (*Right*) Result of aligning λP_R_ class I RPo with a high-resolution X-ray structure of RPinit [2.9 Å resolution, PDB 4Q4Z; (32)]. Initiating triphosphate ribonucleotides (dark beige) bound at +1 (*i* NTP = ATP) and +2 (*i+1* NTP = CMPCPP) pair with the corresponding t-strand bases. The backbone and bases from -3 to +3 in RPinit (beige) largely superpose with the equivalent positions in λP_R_ with the exception of G_-1_(t) where a change in base tilt leads to a clash (red circle) with the iNTP (ATP).

Perhaps surprisingly, no positively charged (or negatively) charged groups fall within 4.5 Å (nor within 6 Å) of the phosphate oxygens from T_-8_(t) to C_-3_(t). By contrast, all interactions with the t-strand DNA near the active site (−2 to +2) are with the DNA backbone [with the exception of σ^70^P427 N near O2 of T_+1_(t)]; salt bridges to each of the phosphate oxygens at -2, -1, and +1 constrain the orientation of the bases with respect to the active site Mg^2+^, leaving the bases free for pairing with substrate.

### λP_R_ class I may be an unproductive RPo

The relatively high resolution of the t-strand in the λP_R_ class I complex (Fig. 1*B*, SI Appendix, Figs. S3 and S12*A*; Table S2) obtained in the absence of NTPs led us to examine whether the observed order corresponds to a site that is pre-organized to bind the first two initiating NTPs. To address this question, we took advantage of the high-resolution X-ray structure [2.9 Å, PDB ID 4Q4Z, (32)] of an initiation complex (RPinit) between *Thermus thermophilus* (*Tth*) Eσ^A^ at a downstream fork promoter construct with the same t-strand sequence from -3 to +1 as λP_R_, and the first two initiating NTPs. In RPinit, the +1 and +2 t-strand bases pair with the initiating nucleotide (*i*NTP=ATP) and a nonhydrolyzable nucleotide analog of the *i+1*NTP (CMPCPP), respectively. Despite differences between the two RNAPs, the DNA constructs used to form RPo and the methods used (X-ray, cryo-EM), the t-strand from -3 to +3 largely superposes (Fig. 5*C*), reflecting the evolutionary conservation of this site in multi-subunit RNAP (48, 49). While the path of the DNA backbone and the plane of the bases at all positions are strikingly similar, G_-1_(t) is a notable exception. In RPinit, G_-1_(t) forms an inter-strand stack with the *i*NTP (32). However, the base tilt of G_-1_(t) in λP_R_ differs: modeling predicts that instead of forming a stabilizing interaction, G_-1_(t) would clash with the incoming *i*NTP, suggesting that this form of RPo at λP_R_ may be recalcitrant to *i*NTP binding (see SI Appendix).

Given the persistent production of abortive products at λP_R_ in transcription assays that prevent RNAP rebinding (single round) (25, 51), we hypothesize that a relatively small conformational change in base tilt at -1 may convert class I λP_R_ to a conformer competent to bind the initiating NTP. Such a movement would occur on a time scale faster than the conversion to a productive complex. At λP_R_, changing the *i*[NTP] from 5 μM to 200 μM increased abortive products by ∼20-fold but had no effect on the amount of full-length RNA (51). One interpretation of these results is that unproductive complexes have a lower NTP binding affinity (51, 52). These data are consistent with the requirement of a conformational change to bind the *i*NTP (Fig. 5*C*).

## Discussion

Our findings illuminate the structural strategies that *Eco* RNAP uses to stabilize the transcription bubble and reveal how small differences in DNA sequence and/or discriminator length globally alter RPo structure. Because the next steps of NTP addition largely disrupt these contacts, their quality and extent impacts how quickly and efficiently RNAP breaks its promoter interactions as it converts to a processive elongation complex (53). Promoter escape is a complex function of DNA sequence and other extrinsic variables [cf. (53) and references therein; (13, 54-57)]. However, in general, initiation from promoters that form highly stable RPo is often rate-limited at escape whereas unstable RPo are limited at the steps of DNA binding and opening [cf. (13, 53)]. As a consequence, unstable RPo (e.g. *rrnb* P1, T7A1) typically produce full length transcripts with few abortive products (58, 59) in single round assays. By contrast, highly stable RPo (e.g. λP_R_) produce smaller amounts of full-length products while continuing to synthesize short RNAs (25, 51, 53, 58, 60).

At all promoters, initiation of RNA synthesis drives translocation of the nascent RNA/DNA hybrid and further unwinding of the downstream duplex DNA. Because Eσ^70^ maintains its interactions with the upstream ds/ss fork junction, the transcription bubble progressively enlarges with each NTP addition, a process termed ‘scrunching’ (61-63). Given the volume constraints of the active site cleft, the cost of DNA unwinding and compaction during initiation was proposed to create “stressed” intermediates and that an accumulation of stress could drive promoter escape (25, 53, 57, 64).

Based on the structures described here, the structural barriers to initially “scrunching” the nt-strand seem low as few or no contacts are made from -3 to -1 in either λP_R_ or T7A1 (Fig. 3). At other promoters, the map density for DNA in this region is poor [λP_R-5C_ (Fig. 2); *rrnB* P1 (36); *rspT* P2 (34)], indicating structural heterogeneity. We note that reducing interactions with the nt-strand discriminator region in RPo appears to favor a state where regions of *both* strands in the bubble are dynamic [cf. (18, 19)]. The T7A1 RPo structure suggests that the first translocation step on a six base discriminator promoter extends the nt-strand downstream of -4 into the channel without steric opposition; subsequent steps are proposed to extrude the nt-strand out between the βprotrusion and βlobe (65). While mid-region contacts are minimal on the t-strand compared to those upstream (Fig. 5*A* and 5*B*), the growing RNA-DNA hybrid pushes the t-strand further into the upstream end of the cleft (65). The t-strand “sandwich” at -10/-9 observed for λP_R_ and T7A1 (Fig. 4) and *rrnB* P1 RPo (36) may provide an additional constraint that helps direct the bulge as scrunching in the t-strand is proposed to disrupt regions (β’lid and the σ-finger) that block RNA entry into its exit channel (65).

### Proposed structural differences between productive and moribund RPo

Seminal work by Shimamoto, Hsu, Chamberlin and colleagues discovered that two forms of RPo can be populated at a given promoter: one that ultimately escapes to make full length RNA, and one stuck in iterative rounds of abortive cycling (51, 52, 60). The latter “moribund” complex often backtracks (3’-OH RNA no longer correctly positioned in the active site), forming a dead-end complex in the absence of Gre factors [cf. (53)]. The relative amounts of these functionally distinct complexes are promoter-sequence dependent (53, 58, 60). One of the striking implications of this work is that core promoter sequence fundamentally sets the degree to which the initiation branches *before* NTP binding [cf. (53) and references therein]. Functionally, branching allows additional regulation of transcriptional output independent of the rate-limiting step of initiation (51) and is modulated by the Gre factors *in vivo* [cf. (66)].

While the structural basis of moribund complex has been unknown, the unambiguous finding of two distinct cryo-EM classes at λP_R_ but not at T7A1 or at λP_R-5C_ may provide an answer. Based on studies of λP_R_ by Shimamoto and colleagues (51, 66) and by the Record lab (21, 22), we suggest that class I and class II represent moribund and productive complexes, respectively (Eq. 1, RPo and I3, see SI Appendix). If so, our data indicate that favorable interactions with the discriminator region (e.g. G_-5_(nt)) drive formation of additional barriers to translocation (increased strand and downstream DNA order, βlobe/σ^70^_1.1_ -linker/discriminator contacts, a more tightly closed clamp). We hypothesize that in this form of RPo, the next step of DNA unwinding is strongly disfavored, leaving the RNA/DNA hybrid largely pre-translocated (the newly added nucleotide remains in the active site). The longer in this state, the greater probability that the 3’ RNA hydroxyl disengages, creating a backtracked complex [cf. (67-69)] that either resets by releasing short RNA products or becomes dead-end complex.

### Implications of this study for the regulation of transcription initiation

Stable transcription bubble formation requires establishing RNAP/DNA interactions that disfavor reannealing, rewinding and DNA dissociation. Promoter sequence intrinsically dictates the cost of disrupting base stacking and base pairing and the degree to which unfavorable conformational changes are driven by forming favorable protein interactions. Perhaps not surprisingly, in the most stable RPo studied here (λP_R_), bases on the separated strands are largely stacked (Fig. 2*A*) and DNA (both base and the phosphate backbone) interactions with Eσ^70^ appear to be maximized relative to the faster dissociating RPo (λP_R-5C_, T7A1). What is more remarkable is how the sequence and length of the nt-strand discriminator impact the overall RPo structure. In particular, the promoter-dependent degrees of disorder of the strands in the bubble described here presumably not only affect RPo lifetime but also start site selection and promoter escape.

The recent cryo-EM study of the steps of DNA opening at the *rpsT* P2 promoter revealed that bubble formation is not a simple progression in strand unwinding and base unstacking: DNA disruptions are dynamic and protein-DNA interactions form and unform (35). Chen et al. found that DNA entry into the active site channel is facilitated by unpairing and unstacking the DNA base pair at -12, which then repairs and restacks with -13 in the subsequent intermediate. In addition, T_-9_(t) binds in a pocket on the βprotrusion at an intermediate step, but then leaves the pocket in subsequent intermediates and RPo (35).

Based on these observations and the comparison between λP_R_ class I and II (Figs. 2*A* and 2*B*), we highlight what may have not been fully appreciated before this structural work: in a multistep mechanism (cf. Eq. 1), if base restacking occurs, it provides an enthalpic driving force for conformational rearrangements in that step, even if the overall net cost to RPo is zero. For example, if unwinding the upstream bubble partially (or fully) unstacks bases in the -10 hexamer [e.g. T_-10_(nt)/A_-9_(nt)/A_-8_(nt) and/or T_-9_(t)] as part of entry into the cleft, the cost is reflected in a slower forward rate or faster back rate (depending whether unstacking occurs before or after the transition state) relative to no disruption. Restacking in a later step will then have the opposite effect on the forward and back rates, respectively. If this scenario is applicable, we note that unstacking purines is more costly than unstacking pyrimidines, suggesting that the base sequence may play hidden roles in the individual steps of DNA opening and in the subsequent steps of promoter escape.

## Materials and Methods

Detailed descriptions of Eσ^70^ purification, assembly of Eσ^70^ – promoter DNA complexes, specimen preparation for cryo-EM, cryo-EM data acquisition and processing, model building and refinement, and abortive initiation transcription assays are provided in SI Appendix.

## Data Availability

The cryo-EM density maps have been deposited in the EMDataBank under accession codes EMD-23892 [*Eco* Eσ^70^-λP_R_ class I (RPo)], EMD-23893 [*Eco* Eσ^70^-λP_R_ class II (I3)], EMD-23895 (*Eco* Eσ^70^-λP_R-5C_ RPo), and EMD-23897 (*Eco* Eσ^70^-T7A1 RPo). The atomic coordinates have been deposited in the Protein Data Bank under accession codes 7MKD [*Eco* Eσ^70^-λP_R_ class I (RPo)], 7MKE [*Eco* Eσ^70^-λP_R_ class II (I3)], 7MKI (*Eco* Eσ^70^-λP_R-5C_ RPo), and 7MKJ (*Eco* Eσ^70^-T7A1 RPo).

## Acknowledgments

We thank E.A. Campbell, M. Lilic and members of the Darst-Campbell Laboratory for experimental advice and helpful discussions. R.M.S. thanks M.T. Record, Jr. and former members of the Record lab for many fruitful collaborations, and T.M. Lohman and D. Jensen for stimulating conversations and encouragement. We are grateful to R.L. Gourse and W. Ross for advice, insightful discussions and inspiring work. Some of the work was performed at the Simons Electron Microscopy Center and National Resource for Automated Molecular Microscopy, located at the New York Structural Biology Center, supported by grants from the Simons Foundation (SF349247), New York State Office of Science, Technology and Academic Research, and the NIH National Institute of General Medical Sciences (GM103310), with additional support from the Agouron Institute (F00316). This work was supported by NIH grant R35 GM118130 and Re-Entry Supplement R35 GM118130-04S1 to S.A.D.

