## Supplemental Information for "Structural origins of *Escherichia coli* RNA polymerase open promoter complex stability"

### Supplemental text

#### Results

##### Characterization of *Eco* E$\sigma^{70}$ RPo at $\lambda P_R$, $\lambda P_{R\text{-}5C}$, and T7A1 promoters under cryo-EM solution conditions

Because RPo stability at a given promoter depends strongly on solution conditions [e.g. salt, solute concentrations, temperature (1-3)], we investigated whether the presence of the detergent CHAPSO affected RPo $t_{1/2}$. In the absence of CHAPSO, very long-lived complexes were populated at $\lambda P_R$ [half-life ($t_{1/2}$) ~10 hours] whereas complexes at T7A1 were short-lived ($t_{1/2}$ ~ 4 minutes; Fig. S1 and Table S1), in agreement with previous investigations (cf. (4, 5)]. The single base pair inversion [$G_{-5}$(nt)] to C in $\lambda P_{R\text{-}5C}$] also significantly reduced RPo $t_{1/2}$ (from 10 to 2 hours; Fig. S1). To illustrate how profound these effects are, Fig. S1 compares the fraction of promoter complexes (theta) populated at 10 minutes after adding competitor: while theta $\lambda P_R$ has not changed outside of uncertainty (100% occupancy, or theta ~ 1), very few T7A1 complexes remain. CHAPSO (8 mM) reduced RPo stability to some extent at all promoters (Fig. S1 and Table S1), but its effects on a given promoter are very small compared to the differences between promoters. As a consequence, 8 mM CHAPSO does not significantly alter the relative stabilities of RPo.

##### CHAPSO binding sites in *Eco* E$\sigma^{70}$ RPo

The structures reported here have CHAPSO bound to various regions of E$\sigma^{70}$, as found in previous cryo-EM studies [cf. (6-9)]. In addition to binding to exterior sites on RNAP, two binding sites are observed in the active site cleft near the $\sigma$-finger [$\sigma^{70}{}_{3.2}$; (10)]. $\lambda P_R$ class I, $\lambda P_{R\text{-}5C}$, and T7A1 RPo have one CHAPSO (I) occupying a region near the t-strand in the vicinity of -6 to -4. $\lambda P_R$ class II and the unstable RPo with the ribosomal promoter *rpsT* P2 (7) have a second CHAPSO (II) bound near CHAPSO (I). The second site clashes with the path of the t-strand in $\lambda P_R$ class II at -6 and -7.

The difference in the number of CHAPSO binding sites between $\lambda P_R$ class I and class II is consistent with the finding that CHAPSO has a slightly larger effect on lifetime on $\lambda P_R$ versus T7A1 or $\lambda P_{R\text{-}5C}$ (which only exhibit one RPo class; Fig. 1*B*; Table S2). Mass action presumably pulls the equilibrium at $\lambda P_R$ to the less stable class II (from a complex with one CHAPSO bound to a complex with two), resulting in a 3-fold effect on $k_d$. CHAPSO likely affects RPo lifetime by competing with a subset of stabilizing interactions in the channel as well as likely having different preferential interactions with the distinct chemical groups on E$\sigma^{70}$ and DNA that will also affect $k_d$ (3). However, for the following reasons we argue that CHAPSO does not affect the main results/conclusions of this study: i. the effects of CHAPSO on RPo lifetime are very small and nowhere near as large as the effects of DNA sequence (Fig. S1, Table S1); ii. t-strand disorder in the vicinity of the CHAPSO active channel binding sites has been found in many RPo X-ray crystal structures (in the absence of CHAPSO); iii. the t-strand (and nt-strand) shows different extents of ordering in cryo-EM structures despite the presence of CHAPSO; iv. the conformation of the $\sigma$-finger is not altered by CHAPSO as the backbone conformer in $\lambda P_R$ class I and in T7A1 is largely the same as that seen in high resolution crystal structures of *T. th.* RPo (11) and RPinit (12); and v. the t-strand backbone and bases (-4 to +2) superpose with X-ray structures of RPinit and RPo ($\lambda P_R$ class I, *rpsT* P2, *rrnB* P1), indicating that any effects of CHAPSO on the t-strand are local and do not propagate to loading in the active site.

Instead of significantly altering the overall path of the t-strand in these structures in solution, we propose the occupancy of the CHAPSO II site correlates with the degree of disorder in the t-strand that exists in RPo in solution in the absence of detergent. For unstable RPo, the t-strand is more dynamic, either never occupying the $\sigma$-finger binding site or having such a low occupancy that CHAPSO binds instead. Recently, a cryo-EM structure of the highly unstable RPo at the *rrnB* P1 promoter (13) was determined using CHAPSO (9). The ability to populate RPo at *rrnb* P1 in the presence of CHAPSO is consistent with the proposal that the detergent binding does not significantly perturb the population of complexes that exists in its absence.

**$\lambda P_R$ class I and class II correspond to RPo and I3, respectively**

Kinetic/mechanistic studies revealed that three open complexes form sequentially at $\lambda P_R$: I2 forms transiently after DNA opening and then is rapidly converted to the more stable complexes I3 and RPo (14, 15). Interpretation of the solute and temperature dependencies of the dissociation rate constant led to the proposal that $\lambda P_R$ RPo is stabilized relative to I3 by "tightening" of the RNAP clamp via interactions involving mobile downstream elements [$\beta$'jaw, sequence insertions Si1 and Si3 (16, 17)] and downstream DNA (5, 18-21). Based on multiple lines of evidence, deletion of either the $\beta$'jaw or DNA downstream of +12 was proposed to prevent the conversion of I3 to RPo (19). Because the cryo-EM $\lambda P_R$ structures are in agreement with these proposals, the simplest interpretation is that class I represents RPo and class II is I3.

**Differences in DNA backbone water-accessibility between $\lambda P_R$ class I and class II provide an explanation for the large effects of temperature and solutes on $k_d$**

The conversion from class II (I3) to class I (RPo) buries additional DNA backbone surface. We estimate that constraining the t-strand from -10 to +2 in the cleft and forming additional interactions with the downstream DNA (+14 to +17) in class I reduces the water-accessible surface area of phosphate oxygens ($\Delta ASA_{OP1,OP2}$) by 620 Å$^2$ relative to class II. This structural estimate is consistent with and provides an explanation for the large effects of temperature and solutes on the RPo dissociation rate constant as outlined below.

First, forming RPo from I3 is enthalpically costly (15). Because the transcription bubble is already open in I3 (14), DNA melting is not the structural origin of $\Delta H^o > 0$. Instead, we suggest the enthalpic cost originates from burying negative charge. Positive enthalpy changes characterize processes that dehydrate anionic groups; release of these waters provides an entropic driving force for forming extensive protein-nucleic acid interfaces (22). Second, solutes that are highly excluded from anionic surface [e.g. the osmolyte glycine betaine (GB) or the physiological anion $Glu^-$] exert extraordinary stabilizing effects on RPo at $\lambda P_R$ (21, 23) and other protein-nucleic acid complexes [cf. (24, 25)]. Because exclusion is so unfavorable, these solutes drive any process that reduces anionic surface. The presence of GB [excluded from three layers of water of hydration (26)] decreases the dissociation rate constant $k_d$ decreases by orders of magnitude: the lifetime of $E\sigma^{70}$-$\lambda P_R$ complexes in 0.5 molal GB at 17.1 °C (21) exceeds a day. By contrast, RPo is not populated at equilibrium at 0.5 M KCl at any temperature (15). Analysis of the dependence of RPo half-life on [GB] using the solute partitioning model [SPM; (3)] predicts that $\Delta ASA_{OP1,OP2}$ ~ -570 Å$^2$ (see Methods below). The semi-quantitative agreement between the prediction based on the class I and II structures and the SPM value suggests that ordering the t-strand and clamp tightening dominate the observed effects of GB on $k_d$ (21).

### Materials and methods

#### Protein expression and purification

***Eco* core RNAP** [$\alpha_2\beta\beta$'(His)$_{10}\omega$] was purified largely as described previously (7). A pET-based plasmid overexpressing each subunit of RNAP (full-length $\alpha$, $\beta$, $\omega$) as well as $\beta$'-PPX-His$_{10}$ (PPX; PreScission protease site, LEVLFQGP, Cytiva, Marlborough, MA) was co-transformed with a pACYCDuet-1 plasmid expressing *Eco rpo*Z into *Eco* BL21(DE3). The cells were grown in the presence of 100 µg/mL ampicillin, 34 $\mu$g/mL chloramphenicol, and 0.5 mm $ZnCl_2$ to an $OD_{600}$ of 0.6 in a 37°C shaker. Protein expression was induced with 1 mM IPTG (final concentration) for 4 hours at 30°C. Cells were harvested by centrifugation and resuspended in lysis buffer [50 mM Tris-HCl, pH 8.0, 5% glycerol (v/v), 10 mM DTT, 1 mM PMSF, and 1x protease inhibitor cocktail (Sigma Aldrich)]. After French Press lysis at 4°C, the lysate was centrifuged twice for 30 minutes each. Polyethyleneimine [PEI, 10% (w/v), pH 8.0, Acros Organics - ThermoFisher Scientific, Waltham, MA] was slowly added to the supernatant to a final concentration of ~0.6% (w/v) PEI with continuous stirring. The mixture was stirred at 4°C for an additional 25 min, then centrifuged for 1.5 hours at 4°C. The pellets were washed three times in lysis buffer + 500 mM NaCl. For each wash, the pellets were homogenized then centrifuged again. RNAP was eluted by washing the pellets three times with lysis buffer + 1 M NaCl. The PEI elutions were combined and precipitated with ammonium sulfate overnight. The mixture was centrifuged and the pellets were resuspended in RNAP buffer [20 mM Tris-HCl, pH 8.0, 5% glycerol (v/v), 5 mM DTT] + 1 M NaCl. The mixture was loaded onto three 5 mL HiTrap IMAC HP columns (Cytiva) for a total column volume (CV) of 15 ml. RNAP($\beta$'-PPX-His$_{10}$) was eluted with RNAP buffer + 250 mM imidazole. The eluted RNAP fractions were combined and dialyzed against RNAP buffer + 100 mM NaCl. The sample was then loaded onto a 35 mL Biorex-70 column (Bio-Rad, Hercules, CA), washed with 10 mM Tris-HCl, pH 8.0, 0.1 mM EDTA, 5% glycerol (v/v), 5 mM DTT, in a gradient from 0.2 M to 0.7 M NaCl. The eluted fractions were combined, concentrated by centrifugal filtration, then loaded onto a 320 mL HiLoad 26/600 Superdex 200 column (GE Healthcare) equilibrated in gel filtration buffer [10 mM Tris-HCl, pH 8.0, 0.1 mM EDTA, 0.5 M NaCl, 5% glycerol (v/v), 5 mM DTT]. The eluted RNAP was supplemented with glycerol to 23% (v/v), flash frozen in liquid $N_2$, and stored at -80°C.

***Eco* His$_{10}$-SUMO-σ$^{70}$ expression and purification.** Plasmid encoding *Eco* His$_{10}$-SUMO-σ$^{70}$ was transformed into *Eco* BL21(DE3) by heat shock. The cells were grown in the presence of 50 μg/mL kanamycin to an $OD_{600}$ of 0.4 at 37°C, then the temperature was lowered to 30$^{o}$C. At OD 0.6, protein expression was induced with 1 mM IPTG (final) for 2 hours. Cells were harvested by centrifugation and resuspended in sigma lysis buffer [20 mM Tris-HCl, pH 8.0, 5% glycerol (v/v), 500 mM NaCl, 0.1 mM EDTA, 5 mM imidazole] and flash frozen in liquid $N_2$. Cells were then thawed on ice, 2-mercaptoethanol (BME) and PMSF were added to 0.5 mM and 1 mM, respectively. After French Press lysis at 4$^{o}$C, cell debris was removed by centrifugation. The lysate was loaded onto two 5 mL HiTrap IMAC HP columns (Cytiva) for a total CV of 10 ml. His$_{10}$-SUMO-σ$^{70}$ was eluted at 250 mM imidazole in 20 mM Tris-HCl, pH 8.0, 500 mM NaCl, 0.1 mM EDTA, 5% glycerol (v/v), 0.5 mM BME. Peak fractions were combined, cleaved with Ulp1, and dialyzed against 20 mM Tris-HCl, pH 8.0, 500 mM NaCl, 0.1 mM EDTA, 5% glycerol (v/v), 0.5 mM BME, resulting in a final imidazole concentration of 25 mM. The sample was loaded onto one 5 mL HiTrap IMAC HP column (Cytiva) to remove His$_{10}$-SUMO-tag along with any remaining uncut σ$^{70}$. Tagless σ$^{70}$ was collected in the flowthrough and concentrated by centrifugal filtration. Pooled, cleaved samples were diluted to 200 mM NaCl using buffer A [10 mM Tris-HCl, pH 8.0, 0.1 mM EDTA, 5% glycerol (v/v), 1 mM DTT] and loaded onto three 5 mL HiTrap Heparin columns (Cytiva) equilibrated in buffer A. The σ$^{70}$ was eluted over a NaCl gradient from 200 mM to 1 M. Peak fractions were pooled, concentrated and buffer exchanged into Superdex equilibration buffer [20 mM Tris-HCl, pH 8.0, 0.5 M NaCl, 5% glycerol (v/v), 1 mM DTT] using centrifugal filtration (Amicon Ultra, MilliporeSigma, Burlington, MA). Concentrated sample was then loaded onto a HiLoad 16/60 Superdex 200 size exclusion column (Cytiva); peak fractions of σ$^{70}$ were pooled, supplemented with glycerol to a final concentration of 20% (v/v), flash-frozen in liquid $N_2$, and stored at -80°C.

#### Preparation of promoter DNA for cryo-EM and abortive initiation assays

Nontemplate (top) and template (bottom) DNA strands (Fig. 1*A*) for each promoter were resuspended in annealing buffer (10 mM Tris-HCl, pH 8.0, 50 mM KCl, 0.1 mM EDTA). Equimolar amounts of the strands were mixed (final concentration 50 μM) and put in a 95$^{o}$C heat block for 5-10 minutes. The heat block was then removed to the benchtop where annealed strands slow cooled to room temperature.

#### Preparation of Eσ$^{70}$-promoter DNA complexes for cryo-EM

Eσ$^{70}$ was formed by mixing core RNAP [$\alpha_2\beta\beta'(\text{His})_{10}\omega$] with a 2-fold molar excess of σ$^{70}$. Samples were incubated for 20-30 minutes at 37$^{o}$C. Eσ$^{70}$ was separated from free σ$^{70}$ on a Superose 6 Increase 10/300 GL size exclusion column (Cytiva) in 10 mM Tris-HCl, pH 8.0, 200 mM KCl, 5 mM $MgCl_2$, 5 mM DTT. Peak fractions of the eluted Eσ$^{70}$ were concentrated to ~10-15 mg/mL by centrifugal filtration (Amicon Ultra, MilliporeSigma). Eσ$^{70}$-λ$P_R$ complexes were formed at room temperature by mixing at a 1(Eσ$^{70}$):1.2

(promoter DNA) molar ratio; E$\sigma^{70}$-$\lambda P_{R\text{-}5C}$ and E$\sigma^{70}$-T7A1 complexes were incubated at room temperature at a 1:1 molar ratio. CHAPSO (3-([3-cholamidopropyl]dimethylammonio)-2-hydroxy-1-propanesulfonate), Anatrace, Maumee, OH) was added to the samples to a final concentration of 8 mM (27). The final buffer condition for all the cryo-EM samples was 10 mM Tris-HCl, pH 8.0, 140 mM KCl, 3.6 mM $MgCl_2$, 3.6 mM DTT, 8 mM CHAPSO. C-flat holey carbon grids (CF-1.2/1.3-4Au, Protochips, Morrisville, NC) were glow-discharged using a Solarus Plasma Cleaner (Gatan, Inc., Pleasanton, CA) for 20 sec prior to the application of 3 µL of sample. Using a Vitrobot Mark IV (FEI, Hillsboro, OR), grids were blotted and plunge-froze into liquid ethane with 100% chamber humidity at 22ºC.

#### Cryo-EM data acquisition and processing

Structural biology software was accessed through the SBGrid consortium (28).

All grids were imaged using a using a 300 keV Titan Krios (ThermoFisher Scientific) equipped with a K2 Summit direct electron detector (Gatan, Inc., Pleasanton, CA). Movies were recorded in super resolution mode; dose-fractionated subframes were 2 x 2 binned, gain-normalized, drifted-corrected, summed, and dose-weighted using MotionCor2 (29). Data were processed using RELION (30) and/or cryoSPARC (31) as described below.

**E$\sigma^{70}$-$\lambda P_R$**. The E$\sigma^{70}$-$\lambda P_R$ dataset was collected at the Simons Electron Microscopy Center (SEMC; New York, NY). Images were recorded with Leginon (32) with a super-resolution pixel size of 0.53 Å over a defocus range of -1 µm to -2.4 µm. Movies were collected at 8.8 electrons/physical pixel/second in dose-fractionation mode with subframes of 0.2 sec over a 6 sec exposure (30 frames) to give a total dose of ~50 electrons/physical pixel. Gctf (33) was used for contrast transfer function estimation. Particles were picked from non-dose weighted images (3,507) using Gautomatch (http://www.mrc-lmb.cam.ac.uk/kzhang/); picked particles (546,990) were extracted from the dose-weighted images (34) using a box size of 256 pixels in RELION (30). Particles were first curated using 2D classification (N=50, 477,285 particles) in cryoSPARC (31); 2D classification of a subset of particles (142,865) generated an *ab initio* 3D map (31) used as a template for heterogeneous refinement. After one round of heterogenous classification (477,285 particles, N=2), 119,794 particles were eliminated as “junk” from the dataset. A second round of heterogeneous classification (N=2) yielded two distinct classes. Class I (267,577 particles) and class II (89,914 particles) were nonuniformly refined (31) in cryoSPARC, yielding 3.2 Å and 3.7 Å nominal resolution sharpened maps, respectively. Local resolution calculations were performed using blocres and blocfilt from the Bsoft package (35).

**E$\sigma^{70}$-T7A1**. Grids containing E$\sigma^{70}$-T7A1 complexes were imaged at the SEMC (New York, NY) as described above for E$\sigma^{70}$-$\lambda P_R$ with the following differences. The total dose was ~43 electrons/physical pixel; defocus range was -1.1 µm to -2.4 µm. The contrast transfer function was estimated for each summed image (6765) using CTFFIND4 (36). Particles (557,254; Gautomatch as above) were extracted from dose-weighted images in RELION (30) using a box size of 320 pixels. A 2D classification of a particle subset (208,939) in cryoSPARC was used as input to generate an *ab initio* reconstruction.

Subsequent homogeneous refinement yielded a 3.4 Å nominal resolution map (cryoSPARC (31)); this map was used as a 3D template for a round of 3D classification (N=3) in RELION done with alignment on all extracted particles. Nonparticles (102,049) identified in this round of classification were removed from the dataset. Remaining particles were combined (455,205), 3D autorefined and postprocessed, and then subjected to three rounds of CTF refinement and Bayesian polishing (34) in RELION (30). Polishing improved the map to a nominal resolution of 3.2 Å. Polished particles were curated using 3D classification without alignment (N=3); remaining "good" particles (346,991) were nonuniformly refined (31) in cyroSPARC, yielding a final map at nominal resolution of 2.9 Å. An attempt to resolve the second, distal $\alpha$-CTD by classifying within a mask around the $\sigma^{70}{}_4$/-35 and upstream DNA (3D subtractive classification) did not improve the map density of this region.

**E$\sigma^{70}$-$\lambda$P$_{R-5C}$.** The E$\sigma^{70}$-$\lambda$P$_{R-5C}$ dataset was collected at the Rockefeller University Evelyn Gruss Lipper Cryo-electron Resource Center. Movies were recorded using Serial EM (37) with a super-resolution pixel size of 0.515 Å over a defocus range of -1.1 µm to -2.4 µm. Movies were collected at 8 electrons/physical pixel/second in dose-fractionation mode with subframes of 0.2 sec over a 10 sec exposure (50 frames) to give a total dose of ~75 electrons/physical pixel. The contrast transfer function was estimated for each summed image (4437) using the Patch CTF module in cryoSPARC (31). Particle picking and extraction from dose-weighted images was done using cryoSPARC Blob Picker and Particle Extraction (box size 256 pixels) modules, respectively. A subset of particles (14,174) chosen from 2D classification (N=50) was used to generate three classes of *ab initio* reconstructions (N=3). These initial models (seed 1: complex; seed 2: decoy; seed 3: complex) were used as 3D templates for cryoSPARC heterogeneous refinement (N=6: two complex models, four decoy templates) on all extracted particles (1,510,888). Particles (396,505) curated from two rounds of heterogeneous refinement done in this manner were homogeneously refined in cryoSPARC (nominal resolution 4.0 Å). After importing refinement data into RELION (30) for two successive iterations of CTF refinement, Bayesian polishing, and 3D auto-refinement (34), the nominal map resolution improved to 3.7 Å. A 3D classification (N=6) of the polished particles without alignment was done in RELION to remove any remaining nonparticles; 3D autorefinement and post-processing of the final set of particles (353,304) in RELION yielded a 3D reconstruction with nominal resolution of 3.5 Å.

#### Model building and refinement of cryo-EM structures

To build initial models of the protein component, E$\sigma^{70}$ from PDB ID 4LJZ (38) was manually fit into the cryo-EM density map of $\lambda$P$_R$ -RPo (class I) using Chimera (39) and real-space refined using PHENIX (40). The DNA was mostly built *de novo* based on the density maps with guidance from the *rpsT* P2 RPo structure (PDB 6OUL; (7)). The E$\sigma^{70}$-$\lambda$P$_R$ class I (RPo) model served as the initial model for E$\sigma^{70}$-$\lambda$P$_R$ class II (I3), E$\sigma^{70}$-T7A1, and E$\sigma^{70}$-$\lambda$P$_{R-5C}$ complexes. The proximal $\alpha$CTD bound to upstream DNA resolved in the E$\sigma^{70}$-$\lambda$P$_R$ class I and E$\sigma^{70}$-T7A1 reconstructions was first modeled using the *rpsT* P2-RPo cryo-EM structure [PDB ID 6OUL (7)]. For real-space refinement, rigid body refinement with sixteen manually-defined mobile domains was followed by all-atom and B-factor refinement with Ramachandran and secondary structure restraints. Models were inspected and modified in Coot (41). Alignment shown in Fig. 5*C* was done using conserved domains of RNAP that exhibit minimal conformational changes in the transcription cycle (6).

#### Surface area calculations and model predictions

Water-accessible surface area (ASA) was calculated using SurfaceRacer5.0 (42) with van der Waals radii from Richards (43) and a probe radius of 1.4 Å. Estimates of the burial of phosphate oxygen ASA in converting I3 to RPo were made by subtracting the ASA of the modeled DNA alone (OP1, OP2; t-strand -10 to +2; duplex DNA +14 to +17) from the corresponding ASA in the $E\sigma^{70}$ complex. This value (620 $Å^2$) was then compared to that predicted by the solute partitioning model (SPM). SPM has successfully quantified the effects of solutes (e.g. Hofmeister effects) on biological processes [cf. (26, 44, 45)]. It posits that the dependence of equilibrium or of rate constants on a given solute concentration (e.g. dlnkd/d[GB]) is directly proportional to the change in surface area ($\Delta$ASA) for that process multiplied by a partition coefficient ($\alpha$). Surface area changes are assigned a chemical type ($\Delta ASA_i$ where i is amide or anionic surface, for example) and partition coefficients are similarly divided based on model biological compound data. For dissociation rate constants, $d(\ln k_d/k_d^o)/d[solute] = \Sigma$ (over i) $\alpha_i \Delta ASA_i$ (where $k_d^o$ is the half-life at 0 molal GB). For the interaction of GB with phosphate oxygens, $\alpha_i = 4.9 \times 10^{-3}$ $m^{-1}$ $Å^{-2}$ (26). Based on the data of Kontur et al. (2006)(21), $d\ln(t_{1/2}/t\ t_{1/2}^o)/d[GB] = 2.8\ m^{-1}$. [RPo complexes are so long-lived in GB that empirical half-lives were used to describe the effects of GB. Up to ~20 hours, dissociation data (60-70%) are well-described by a single exponential; afterwards the decay is slower than expected (21).]

#### Promoter lifetime and abortive initiation transcription assays

RPo lifetimes were determined in the presence and absence of 8 mM CHAPSO using the abortive initiation assay (46). Because heparin does not act as inert competitor in measuring the dissociation of promoter complexes formed at T7A1 (4), a consensus "bubble" promoter construct was used to trap $E\sigma^{70}$ irreversibly after it dissociated [bubble promoter DNA: consensus UP element (47), -35 hexamer (TTGACA), "extended" -10 TG at positions -15, -14 (48), -10 hexamer (TATACT), discriminator GGG -6 to -4 (49) and an 11 base mismatched bubble from -11 to +1]. RPo stability at T7A1 and at $\lambda P_{R\text{-}5C}$ was determined at 37°C in transcription buffer (TB: 40 mM Tris-HCl, pH 8.0, 120 mM KCl, 10 mM $MgCl_2$, 100 μg/mL BSA, and 1 mM DTT). To bring the dissociation rate of $\lambda P_R$ RPo complexes into a more experimentally-accessible time range [$t_{1/2}$ RPo at $\lambda P_R$ in TB at 37°C exceeds 8 hours; (15, 50)], the effects of CHAPSO were determined at lower temperature (24°C) and higher [KCl] (8 mM Tris HCl, pH 8.0, 160 mM KCl, 8 mM $MgCl_2$, 100 μg/mL BSA, 1 mM DTT). For all assays, $E\sigma^{70}$ (20 nM or 40 nM final) and promoter DNA (40 nM or 86 nM final) were incubated for 30-60 minutes. After addition of competitor (t=0), samples were removed at time t and allowed to transcribe for 10 minutes in the presence of 100 μM CpA (TriLink Biotechnologies, San Diego, CA), 10 μM UTP (Cytiva) and 1-2 μCi of [$\alpha$-$^{32}$P]-UTP (Perkin Elmer Life

Sciences, Waltham, MA). Labeled trinucleotides were separated from unincorporated [$\alpha$-$^{32}$P]-UTP on 20% polyacrylamide, 6 M urea gels. Gels were imaged on a Typhoon phosphorimager (Cytiva). Trinucleotide bands were boxed, quantified and background corrected using ImageJ (51). Dissociation rate constants ($k_d$) were determined using Prism 7.0 (GraphPad Software, Inc., San Diego, CA). Table S1 presents the mean values and the standard error of the mean from three independent determinations.

#### Supplemental Figures

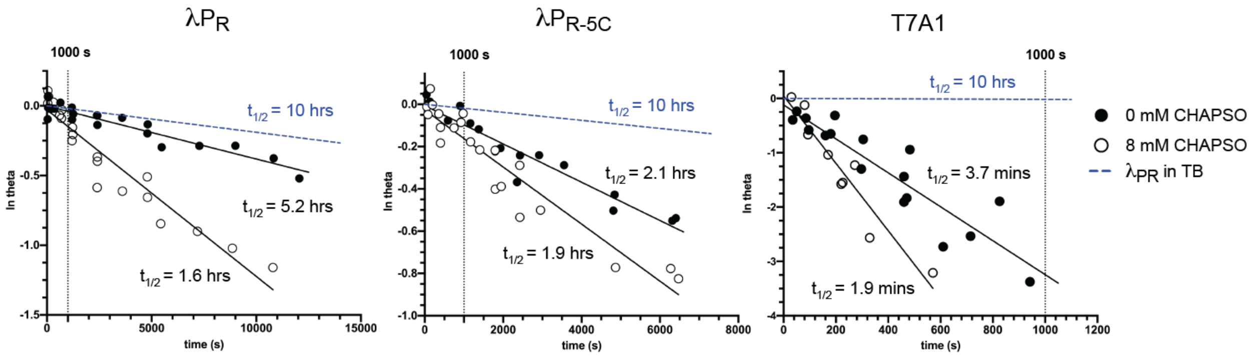

**Fig. S1.** Comparison of the RPo lifetime as a function of DNA sequence in the presence and absence of 8 mM CHAPSO. Promoter occupancy of RPo (theta) at $\lambda P_R$, $\lambda P_{R\text{-}5C}$ or T7A1 was measured as a function of time the after addition of an inert competitor in the absence (open circle) and presence (filled circle) of the detergent used in cryo-EM grid preparation (8 mM CHAPSO). $\lambda P_{R\text{-}5C}$ and T7A1 were investigated in TB buffer at 37$^o$C. Because the half-life of RPo at $\lambda P_R$ in TB at 37$^o$C is ~10 hours, the effect of 8 mM CHAPSO on $\lambda P_R$ dissociation was determined using a higher salt buffer (destabilizing) and a lower temperature (24$^o$C). To facilitate comparison of the relative stabilities of RPo, each plot includes: i. a blue dashed line representing the dissociation of RPo at $\lambda P_R$ at 37$^o$C in TB buffer ($k_d$ = 1.9 x 10$^{-5}$ s$^{-1}$, $t_{1/2}$ = 10 hours; (15); ii. a black dotted line highlighting a common time point (600 s). See Table S1 for $k_d$ values and half-lives.

**Fig S2.** Cryo-EM processing pipeline for $E\sigma^{70}$ - $\lambda P_R$ promoter DNA complexes.

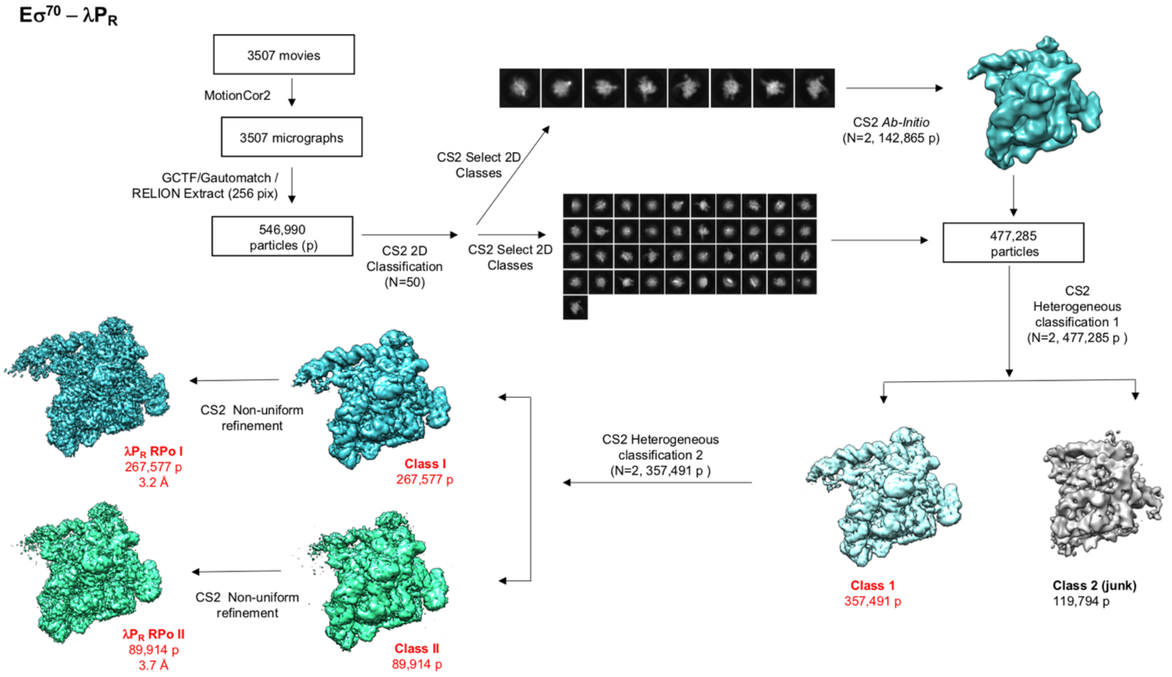

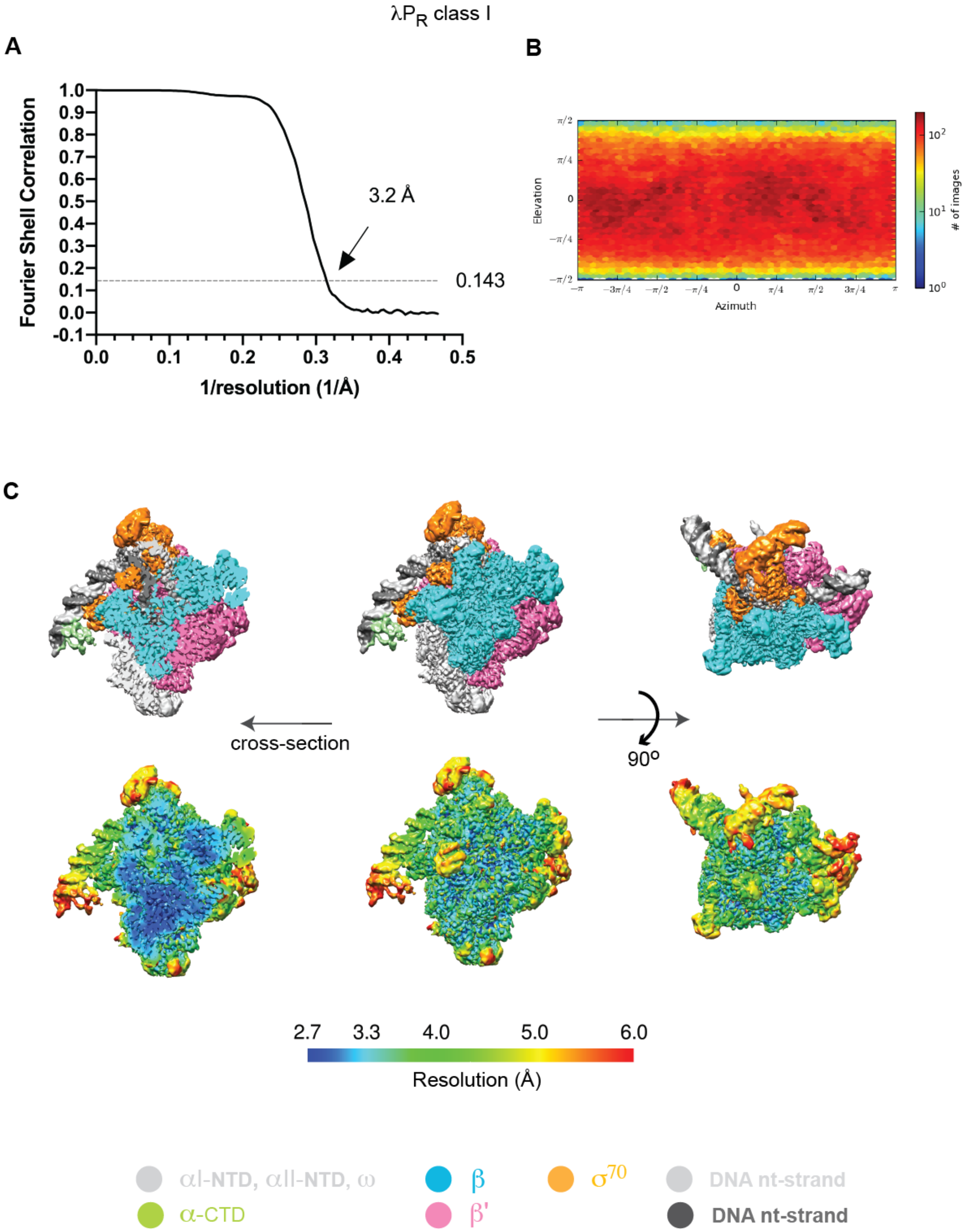

**Fig. S3.** Cryo-EM of E$\sigma^{70}$-$\lambda P_R$ class I.

(**A**) Gold standard FSC plot (52) calculated by comparing the two independently-determined half maps from cryoSPARC (31). Nominal resolution values for sharpened map shown at the 0.143 FSC cutoff.

(**B**) 3D FSC plot (53). Resolution corresponds to the unsharpened map.

(**C**) Heat map showing the angular distribution of particle views calculated in cryoSPARC.

(**D**) Different representations and views of the cryo-EM maps for RPo formed at each promoter.

(*Top*) Cryo-EM density map shown in different views; subunits and DNA strands colored as in Fig. 1.

(*Bottom*) Local resolution of the cryo-EM density map shown above calculated using Bsoft (35), resolution coloring key shown directly below.

(*Left*) Cross-section of the view shown in the (*Middle*) showing the path and resolution of the nt- and t-strands and the relatively high resolution (blue) of the RNAP "core" and active site channel.

(*Right*) ~90$^{o}$ rotation around the x-axis of middle view, chosen to show i. the enclosure of the transcription bubble by a "clasp" formed between the $\beta$'clamp and $\beta$lobe; ii. the extent of cryo-EM density for the N-terminal residues of $\sigma^{70}$ interacting with the $\beta$ lobe; and iii. the extent of cryo-EM density for the downstream DNA.

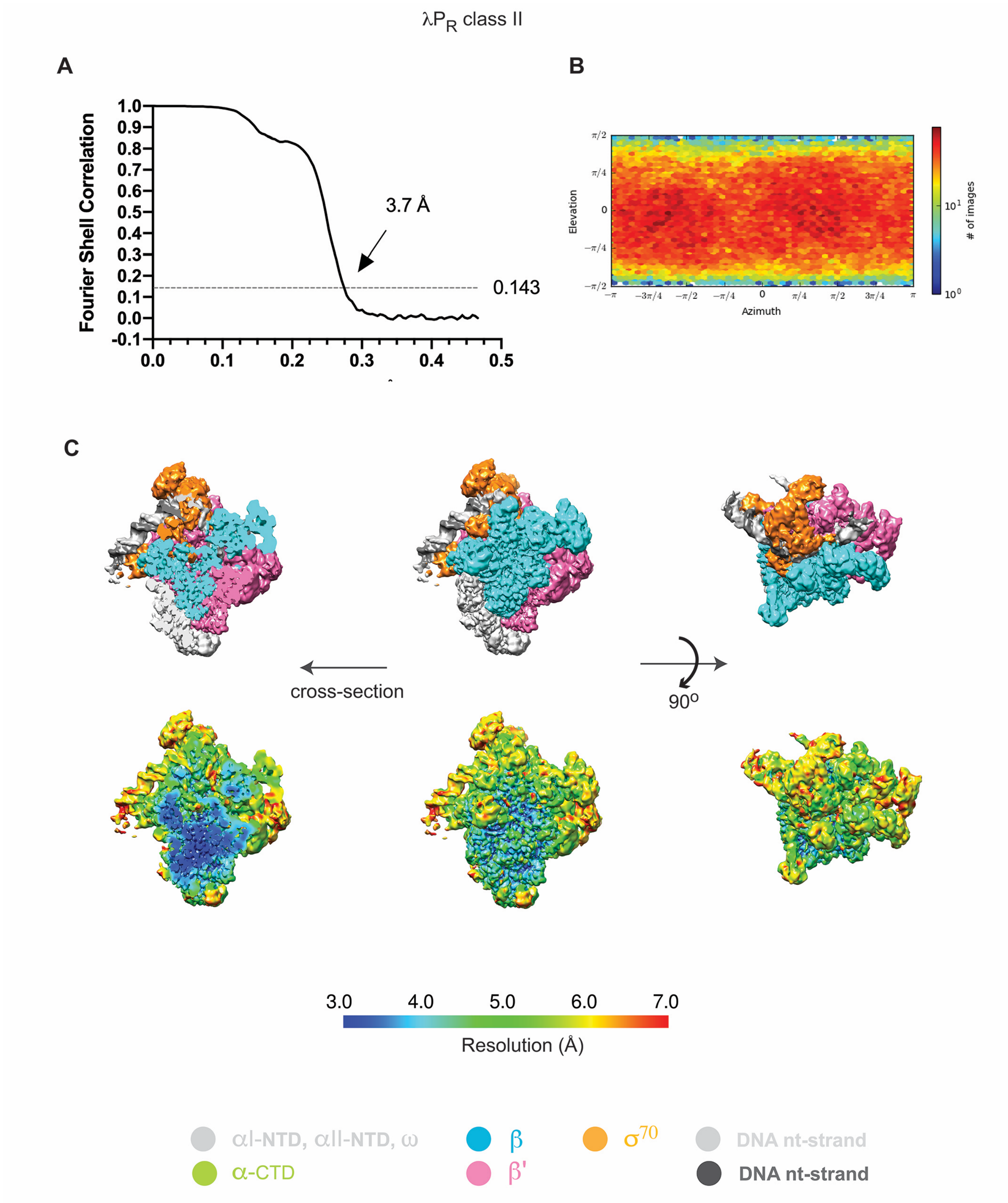

**Fig. S4.** Cryo-EM of E$\sigma^{70}$-$\lambda P_R$ class II.

(**A**) Gold standard FSC plot (52) calculated by comparing the two independently-determined half maps from cryoSPARC (31). Nominal resolution values for sharpened map shown at the 0.143 FSC cutoff.

(**B**) 3D FSC plot (53). Resolution corresponds to the unsharpened map.

(**C**) Heat map showing the angular distribution of particle views calculated in cryoSPARC.

(**D**) Different representations and views of the cryo-EM maps for RPo formed at each promoter.

(*Top*) Cryo-EM density map shown in different views; subunits and DNA strands colored as in Fig. 1.

(*Bottom*) Local resolution of the cryo-EM density map shown above calculated using Bsoft (35), resolution coloring key shown directly below.

(*Left*) Cross-section of the view shown in the (*Middle*) showing the path and resolution of the nt- and t-strands and the relatively high resolution (blue) of the RNAP "core" and active site channel.

(*Right*) ~90$^{o}$ rotation around the x-axis of middle view, chosen to show i. the enclosure of the transcription bubble by a "clasp" formed between the $\beta$'clamp and $\beta$lobe; ii. the extent of cryo-EM density for the N-terminal residues of $\sigma^{70}$ interacting with the $\beta$ lobe; and iii. the extent of cryo-EM density for the downstream DNA.

**Fig. S5.** Cryo-EM processing pipeline for E$\sigma^{70}$-$\lambda P_{R\text{-}5C}$ promoter DNA complexes.

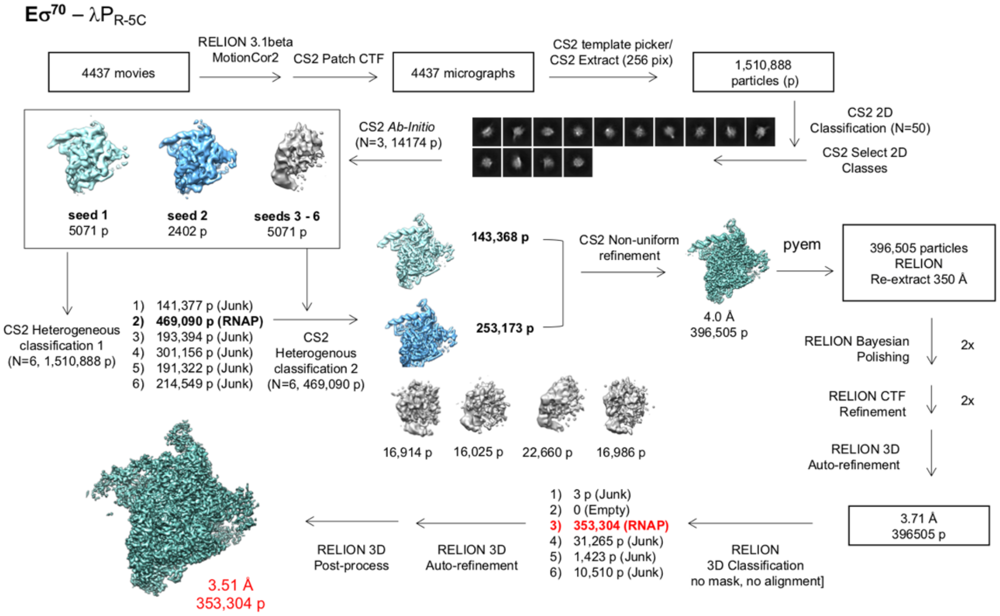

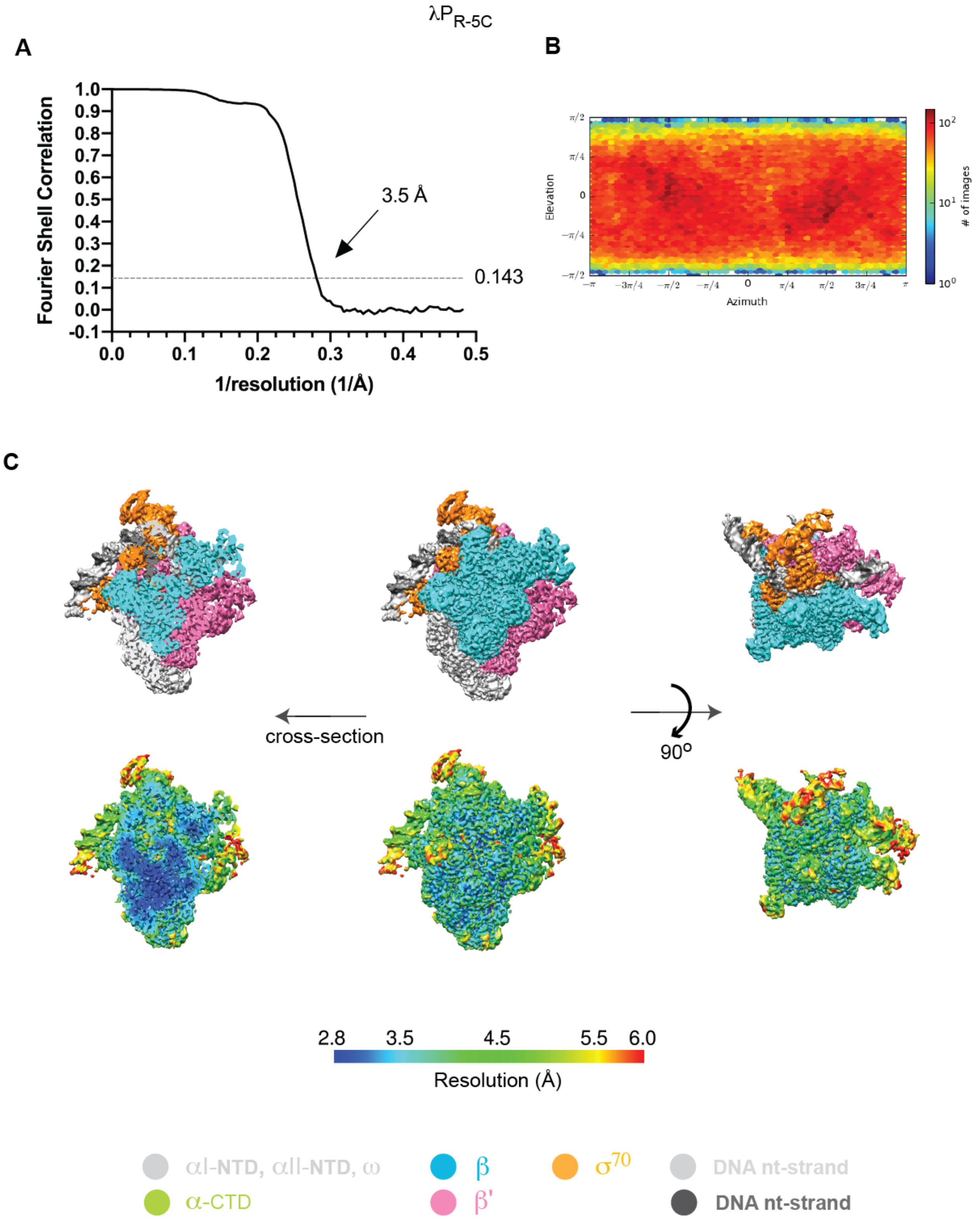

**Fig. S6.** Cryo-EM of E$\sigma^{70}$-$\lambda P_{R\text{-}5C}$.

(**A**) Gold standard FSC plot (52) calculated by comparing the two independently-determined half maps from cryoSPARC (31). Nominal resolution values for sharpened map shown at the 0.143 FSC cutoff.

(**B**) 3D FSC plot (53). Resolution corresponds to the unsharpened map.

(**C**) Heat map showing the angular distribution of particle views calculated in cryoSPARC.

(**D**) Different representations and views of the cryo-EM maps for RPo formed at each promoter.

(*Top*) Cryo-EM density map shown in different views; subunits and DNA strands colored as in Fig. 1.

(*Bottom*) Local resolution of the cryo-EM density map shown above calculated using Bsoft (35), resolution coloring key shown directly below.

(*Left*) Cross-section of the view shown in the (*Middle*) showing the path and resolution of the nt- and t-strands and the relatively high resolution (blue) of the RNAP "core" and active site channel.

(*Right*) ~90$^{o}$ rotation around the x-axis of middle view, chosen to show i. the enclosure of the transcription bubble by a "clasp" formed between the $\beta$'clamp and $\beta$lobe; ii. the extent of cryo-EM density for the N-terminal residues of $\sigma^{70}$ interacting with the $\beta$ lobe; and iii. the extent of cryo-EM density for the downstream DNA.

**Fig. S7.** E$\sigma^{70}$ cryo-EM processing pipeline for E$\sigma^{70}$-T7A1 promoter DNA complexes.

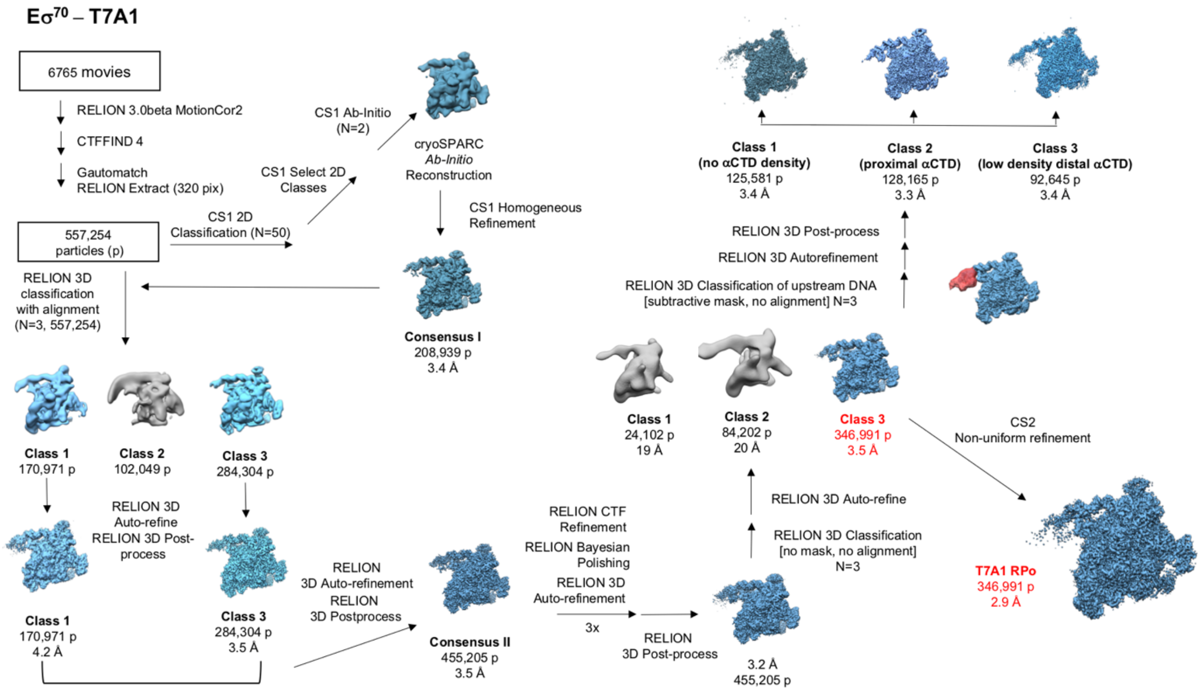

**T7A1**

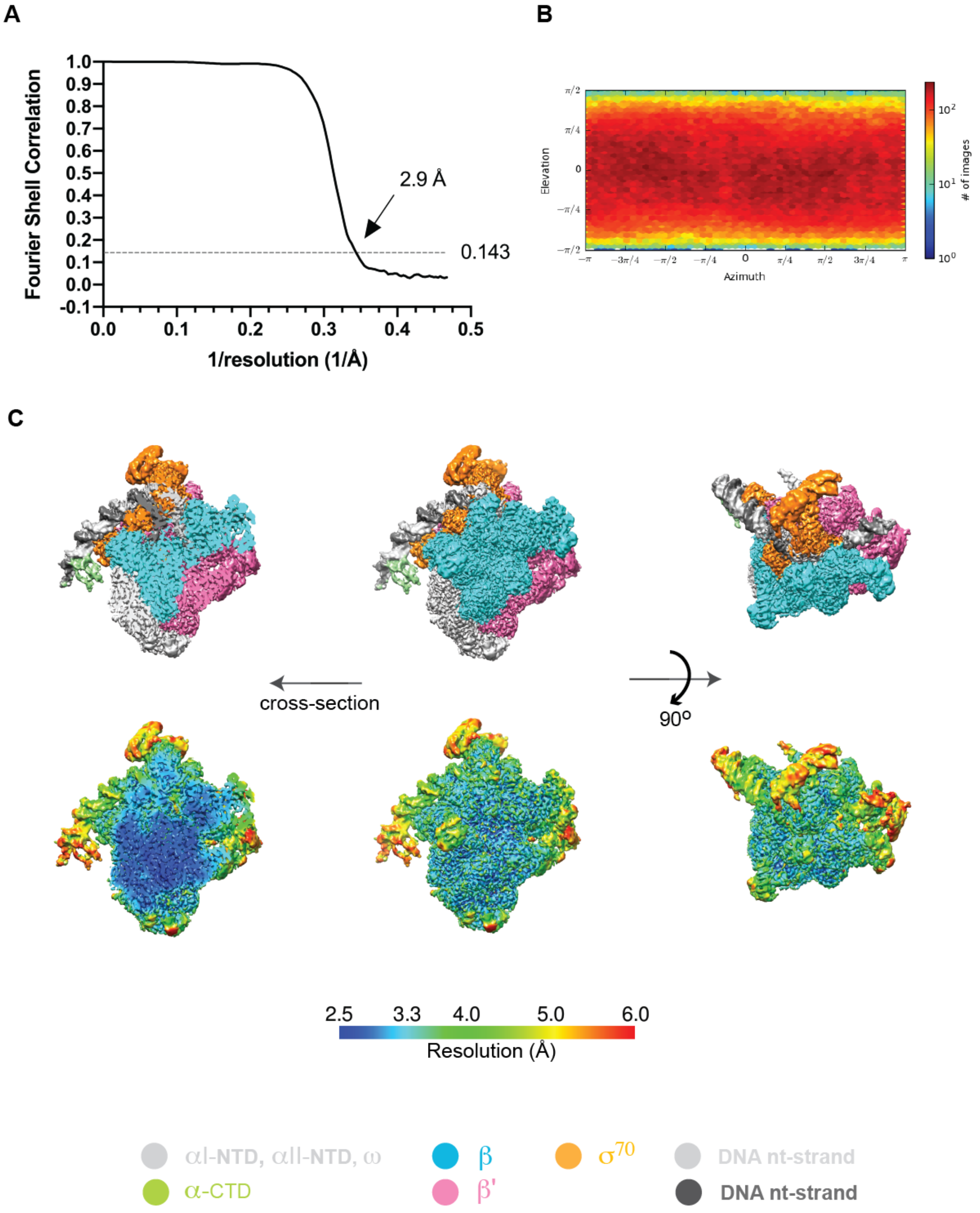

**Fig. S8.** Cryo-EM of E$\sigma^{70}$-T7A1.

(**A**) Gold standard FSC plot (52) calculated by comparing the two independently-determined half maps from cryoSPARC (31). Nominal resolution values for sharpened map shown at the 0.143 FSC cutoff.

(**B**) 3D FSC plot (53). Resolution corresponds to the unsharpened map.

(**C**) Heat map showing the angular distribution of particle views calculated in cryoSPARC.

(**D**) Different representations and views of the cryo-EM maps for RPo formed at each promoter.

(*Top*) Cryo-EM density map shown in different views; subunits and DNA strands colored as in Fig. 1.

(*Bottom*) Local resolution of the cryo-EM density map shown above calculated using Bsoft (35), resolution coloring key shown directly below.

(*Left*) Cross-section of the view shown in the (*Middle*) showing the path and resolution of the nt- and t-strands and the relatively high resolution (blue) of the RNAP "core" and active site channel.

(*Right*) ~90° rotation around the x-axis of middle view, chosen to show i. the enclosure of the transcription bubble by a "clasp" formed between the β'clamp and βlobe; ii. the extent of cryo-EM density for the N-terminal residues of $\sigma^{70}$ interacting with the β lobe; and iii. the extent of cryo-EM density for the downstream DNA.

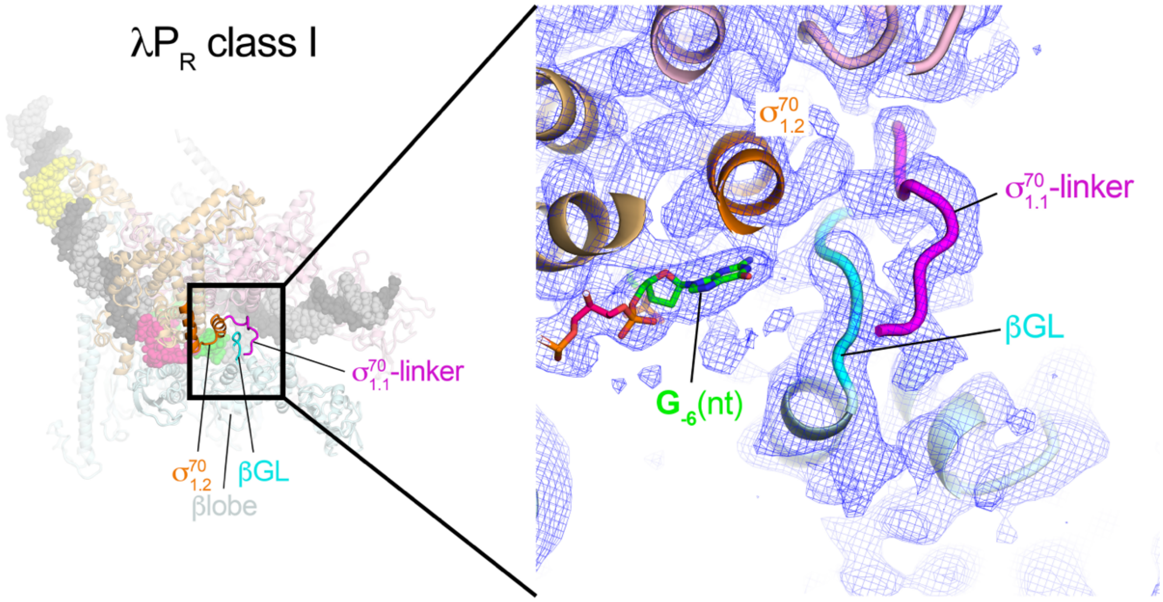

**Fig. S9. Extra residues of σ$^{70}$$_{1.1}$-linker ordered in λP$_R$ class I.**

(*left*) View of overall λP$_R$ class I structure. The boxed region is magnified on the right.

(*right*) Magnified view showing proteins as backbone worms. The cryo-EM density (local-resolution filtered) is shown as a blue mesh.

In class λP$_R$ II, the relatively open state of the clamp (Table S3) positions G$_{-6}$(nt) and G$_{-5}$(nt) away from the βGL and σ$^{70}$$_{1.2}$ such that the key residues (βR371, σ$^{70}$D96, σ$^{70}$R99, σ$^{70}$M102; see Fig. 3*A*) cannot form favorable interactions. Closure of the "clasp" in class I (RPo) not only creates the extensive network between σ$^{70}$$_{1.2}$, σ$^{70}$$_2$, and the βGL, it also allows the formation of additional interactions between σ$^{70}$ residues 85–89 and the βlobe. These contacts may direct the negatively charged σ$^{70}$$_{1.1}$ away from the active site cleft. Because this interaction is only seen for λP$_R$ RPo, it seems likely that discriminator length and sequence determine the degree to which interactions between the βGL and σ$^{70}$$_{1.2}$ are favorable enough to drive additional folding of the σ$^{70}$$_{1.1}$-linker. While the βGL and σ$^{70}$$_{1.2}$ form a clasp in all the RPo reported here, the remaining σ$^{70}$$_{1.1}$-linker can adopt two states. In state 1, residues immediately N-terminal to σ$^{70}$V94 remain disordered, resulting in an ensemble of states where the local concentration of σ$^{70}$$_{1.1}$ remains relatively high above the cleft [λP$_R$ class II, T7A1, λP$_{R\text{-}5C}$, *rpsT* P2 (PDB 6OUL; (7), *rrnB* P1 (PDB 7KBH; (9)]. In state 2, ordering and binding of amino acids N terminal to 94 to the top of the βlobe, direct σ$^{70}$$_{1.1}$ away from the cleft (λP$_R$ class I). In the more "locked" state 2, the local concentration of σ$^{70}$$_{1.1}$ vis a vis the channel is reduced, lowering the probability of σ$^{70}$$_{1.1}$ reinvading the cleft and displacing DNA.

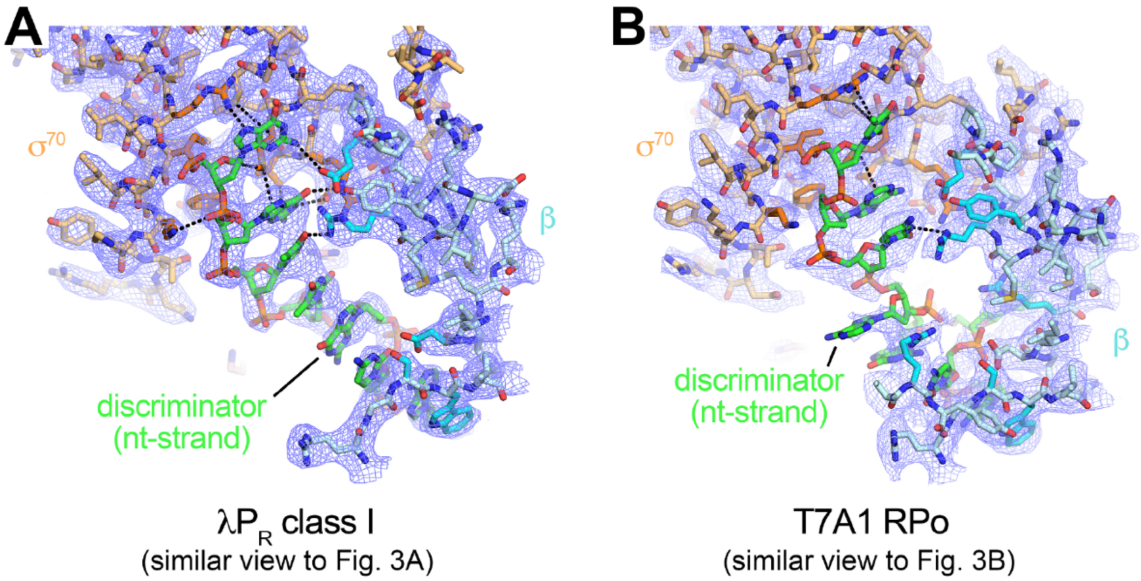

**Fig. S10**. Cryo-EM density for the nt-strand discriminator regions of (**A**) $\lambda P_R$ class **I**, and (**B**) T7A1. Views are very similar to the views of Fig. 3.

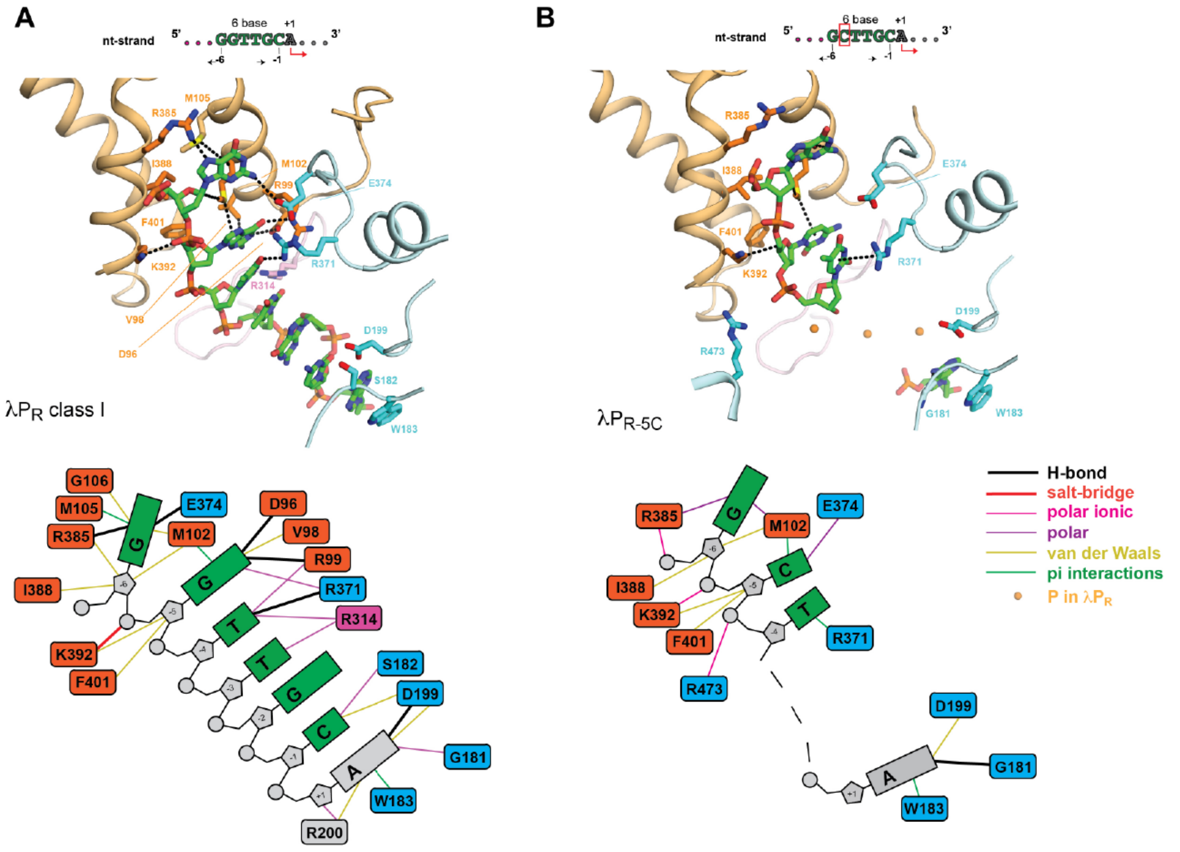

**Fig. S11**. Differences in E$\sigma^{70}$ interactions with the nt-strand discriminator region between (**A**) $\lambda P_R$ class I and (**B**) $\lambda P_{R\text{-}5C}$.

Even though $\lambda P_R$ and $\lambda P_{R\text{-}5C}$ have identical bases at $G_{-6}$(nt) and $T_{-4}$(nt), all hydrogen bonds with these bases are lost in $\lambda P_{R\text{-}5C}$. Repositioning of the clamp (more open; Table S3) and changes in the path of the nt-strand downstream of $T_{-7}$(nt) eliminate the sulfur-pi interaction between $G_{-6}$(nt) and $\sigma^{70}$M105 and the van der Waals contacts with $\sigma^{70}$G106. In addition, conserved residues $\sigma^{70}$D96, $\sigma^{70}$V98 and $\sigma^{70}$R99 no longer contact these bases in $\lambda P_{R\text{-}5C}$. (*Top*) Interactions with the nt-strand from -6 to +1. In $\lambda P_{R\text{-}5C}$, the nt-strand DNA from -3 to -1 lacks interpretable cryo-EM density; orange-yellow spheres are positioned approximately at the corresponding positions of the phosphate backbone in $\lambda P_R$ class I to represent the covalent connection from -4 to +1. E$\sigma^{70}$ subunits shown in backbone worm; sidechains of atoms within 4.5 Å of nucleic acid atoms are shown as sticks. Atomic distances within 3.5 Å and interactions with the pi electrons of the DNA bases (sulfur-$\pi$, and cation-$\pi$) are shown by black dashed lines (coloring scheme as in Fig. 3). (*Bottom*) Schematic comparing the network of interactions between the nt-discriminator (and +1) and E$\sigma^{70}$. Favorable interactions that are within 3.5 Å (hydrogen bonds, salt bridges) are shown by heavier lines (see color key) than those within 4.5 Å (polar, ionic, van der Waals, pi (sulfur, cation, $\pi$)). Bases -3 to -1 in $\lambda P_{R\text{-}5C}$ are shown as black dashes.

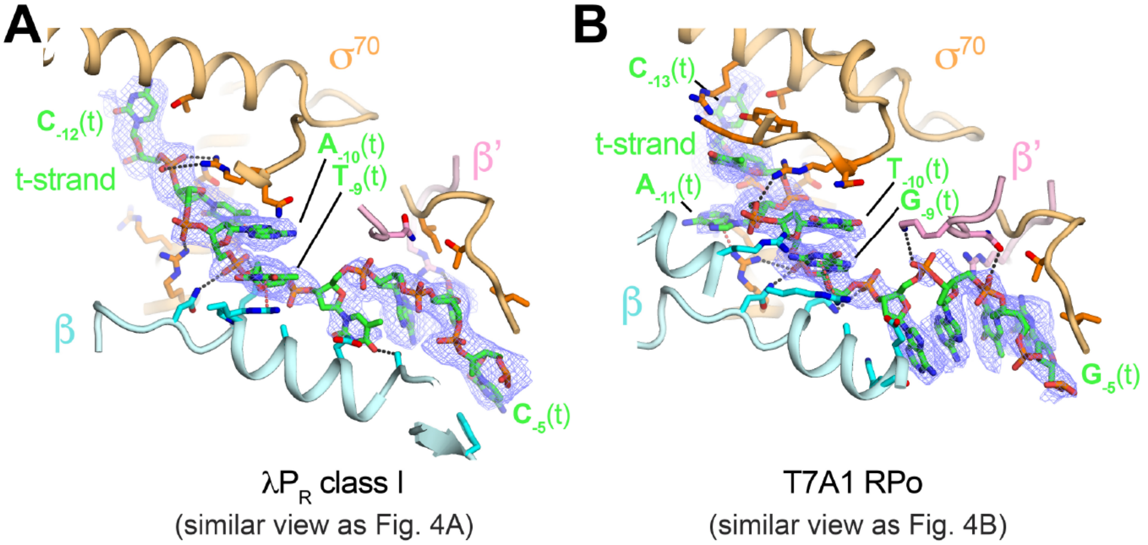

**Fig. S12**. Cryo-EM density (blue mesh, for DNA only) for the t-strand 'sandwich' regions of (**A**) λ$P_R$ class **I**, and (**B**) T7A1. Views are very similar to the views of Fig. 4.

#### Supplemental Tables

**Table S1**. Dissociation rate constants and half-lives ($t_{1/2}$) as a function of promoter and of CHAPSO.

| Promoter | 0 mM CHAPSO | | 8 mM CHAPSO | | Fold-difference[a] |
|---|---|---|---|---|---|
| | $k_d$ ($s^{-1}$) | $t_{1/2}$ | $k_d$ ($s^{-1}$) | $t_{1/2}$ | |
| $\lambda_{PR}$ [b] | $(3.8 \pm 0.6) \times 10^{-5}$ | 5.2 hours | $(1.2 \pm 0.2) \times 10^{-4}$ | 1.6 hours | 3.2 |
| $\lambda_{PR}$ [c] | $(1.9 \pm 0.6) \times 10^{-5}$ | 10.1 hours | N.D. | N.D. | |
| $\lambda_{PR\text{-}5C}$ | $(9.1 \pm 0.6) \times 10^{-5}$ | 2.1 hours | $(1.3 \pm 0.2) \times 10^{-4}$ | 1.9 hours | 1.4 |
| T7A1 | $(3.1 \pm 0.4) \times 10^{-3}$ | 3.7 minutes | $(6.2 \pm 0.8) \times 10^{-3}$ | 1.9 minutes | 2.0 |

a. $k_d$ CHAPSO (8 mM) / $k_d$ CHAPSO (0 mM).

b. Dissociation was measured in buffer containing 8 mM Tris HCl, pH 8.0, 160 mM KCl, 8 mM $MgCl_2$, 100 μg/mL BSA, 1 mM DTT at 24°C.

c. Value of $k_d$ for $\lambda P_R$ determined under the same solution conditions as the measurements for $\lambda_{PR\text{-}5C}$ and T7A1 (37°C, 120 mM KCl, 10 mM $MgCl_2$, 40 mM Tris pH 8.0, 100 μg/mL BSA, 1 mM DTT; (15)).

**Table S2.** Cryo-EM data acquisition and refinement parameters.

| **Sample** | $E\sigma^{70}$-$\lambda P_R$ | | $E\sigma^{70}$-$\lambda P_{R\text{-}5C}$ | $E\sigma^{70}$-T7A1 |
|---|---|---|---|---|
| **Data collection and processing** | | | | |
| Microscope (ThermoFisher Scientific) | Titan Krios | | Titan Krios | Titan Krios |
| Voltage (kV) | 300 | | 300 | 300 |
| Detector (Gatan) | K2 summit | | K2 summit | K2 summit |
| Electron exposure ($e^{-}$/Å$^2$) | 46 | | 75 | 43 |
| Defocus range (μm) | -1.0 to -2.2 | | -1.1 to -2.4 | -1.1 to -2.4 |
| Data collection mode | Super-resolution | | Super-resolution | Super-resolution |
| Physical pixel size (Å) | 1.06 | | 1.03 | 1.06 |
| Symmetry imposed | C1 | | C1 | C1 |
| Initial particle images (no.) | 546,990 | | 1,510,888 | 557,254 |
| Final particle images (no.) | 357,491 | | 353,304 | 346,991 |
| | Class I | Class II | | |
| EMDB | EMD-23892 | EMD-23893 | EMD-23895 | EMD-238957 |
| PDB | 7MKD | 7MKE | 7MKI | 7MKJ |
| final particle images (no) | 267,577 | 89,914 | 353,304 | 346,991 |
| Map resolution (Å) - FSC threshold 0.143 | 3.2 | 3.7 | 3.5 | 2.9 |
| Map resolution range (Å) | 2.7 - 6.0 | 3.0 - 7.0 | 2.8 - 6.0 | 2.5 - 6.0 |
| **Refinement**[c] | | | | |
| Initial model used (PDB code) | 4LJZ[a]/6OUL[b] | 4LJZ[a]/6OUL[b] | 4LJZ[a]/6OUL[b] | 4LJZ[a]/6OUL[b] |
| Map sharpening B factor (Å$^2$) | 133.0 | 109.0 | -121.3 | 107.1 |
| Model composition | | | | |
| Non-hydrogen atoms | 31,518 | 30,785 | 30,979 | 32,095 |
| Protein residues | 3,668 | 3,682 | 3,683 | 3,754 |
| Nucleic acid residues | 120 | 85 | 96 | 122 |
| Ligands | 1 $Mg^{2+}$,<br>2 $Zn^{2+}$,<br>3 CHAPSO | 1 $Mg^{2+}$,<br>2 $Zn^{2+}$,<br>4 CHAPSO | 1 $Mg^{2+}$<br>2 $Zn^{2+}$<br>3 CHAPSO | 1 $Mg^{2+}$<br>2 $Zn^{2+}$<br>3 CHAPSO |
| B factors (Å$^2$) | | | | |
| Protein | 112.00 | 117.34 | 58.96 | 70.76 |
| Nucleic acid | 175.45 | 253.65 | 145.09 | 196.64 |
| Ligands | 101.89 | 94.05 | 54.79 | 64.93 |
| R.m.s. deviations | | | | |
| Bond lengths (Å) | 0.008 | 0.008 | 0.006 | 0.005 |
| Bond angles (°) | 0.760 | 0.838 | 0.735 | 0.645 |
| Validation | | | | |

| | | | | |
|---|---|---|---|---|
| Clashscore | 6.69 | 8.95 | 6.39 | 5.49 |
| Poor rotamers (%) | 5.35 | 7.34 | 5.21 | 3.79 |
| Ramachandran plot | | | | |
| Favored (%) | 93.32 | 90.99 | 92.57 | 94.61 |
| Allowed (%) | 6.62 | 8.96 | 7.40 | 5.39 |
| Disallowed (%) | 0.05 | 0.05 | 0.03 | 0.0 |

[a](38).

[b](7).

[c]Refinement: PHENIX real_space_refine (40). Validation: MolProbity (54).

**Table S3. Rotation analysis of the clamp and the βlobe**

| Eσ$^{70}$ RPo | λP$_R$ Class I reference | |
|---|---|---|
| | clamp[a] | βlobe[b] |
| λP$_R$ class I | (0$^o$) | (0$^o$) |
| λP$_R$ class II | 1.66$^o$ | 1.29$^o$ |
| λP$_{R\text{-}5C}$ | 0.66$^o$ | 0.86$^o$ |
| T7A1 | 0.92$^o$ | 1.29$^o$ |
| *rpsT* P2 (PDB ID 6OUL)[c] | 0.94$^o$ | 1.38$^o$ |
| *rrnB* P1 (PDB ID 7KHB)[d] | 0.73$^o$ [e] | 1.26$^o$ |

[a]Clamp residues: β1319-1342; β'1-342, β'318-1344, σ$^{70}$ 90-137, and σ$^{70}$353-449.
[b]βlobe residues: β152-443.
[c](7).
[d](9).
[e]The axis of rotation for *rrnB* P1 with respect to λP$_R$ class I is not open-close with respect to the upstream/downstream axis of the channel but instead is more swivel-like (6), twisting with respect to the clamp position in λP$_R$ class I. *rrnB* P1 has a short 16 base pair spacer. Promoters with spacers longer than the consensus 17 base pairs kink the DNA in between the -35 and -10 element to maintain interactions with σ$^{70}{}_2$ and σ$^{70}{}_4$, respectively. The swivel in the clamp may arise to accommodate the shorter nonconsensus 16 base pair spacer [cf. (55)].
